# Transcriptional signatures of cell-cell interactions are dependent on cellular context

**DOI:** 10.1101/2021.09.06.459134

**Authors:** Brendan T. Innes, Gary D. Bader

## Abstract

Cell-cell interactions are often predicted from single-cell transcriptomics data based on observing receptor and corresponding ligand transcripts in cells. These predictions could theoretically be improved by inspecting the transcriptome of the receptor cell for evidence of gene expression changes in response to the ligand. It is commonly expected that a given receptor, in response to ligand activation, will have a characteristic downstream gene expression signature. However, this assumption has not been well tested. We used ligand perturbation data from both the high-throughput Connectivity Map resource and published transcriptomic assays of cell lines and purified cell populations to determine whether ligand signals have unique and generalizable transcriptional signatures across biological conditions. Most of the receptors we analyzed did not have such characteristic gene expression signatures – instead these signatures were highly dependent on cell type. Cell context is thus important when considering transcriptomic evidence of ligand signaling, which makes it challenging to build generalizable ligand-receptor interaction signatures to improve cell-cell interaction predictions.

## Introduction

The advent of single-cell RNA sequencing has enabled researchers to explore the cell composition of complex tissues and better understand how cells work together in the context of the entire tissue. Cell-cell interaction prediction from scRNAseq data is useful to support this ‘cellular ecosystem’ study ((Shao *et al*, 2020), (Armingol *et al*, 2021), see Supplementary Table 1 for a comprehensive summary of published methods). The fundamental idea behind cell-cell interaction inference is that if one cell type expresses a ligand, and another cell type expresses its cognate receptor, these cell types may be communicating via paracrine signaling through this putative ligand-receptor interaction. Given an adequate database of ligand-receptor interacting pairs, one can quickly generate lists of candidate interactions between cell types, represented by scRNAseq-derived cell clusters. The disadvantage is that by ignoring myriad factors not directly assayed by scRNAseq that affect ligand-receptor signaling (e.g. post-translational regulation, protein expression level, cellular localization, small molecule cofactors, receptor dimerization), these predictions lack specificity. Most existing cell-cell interaction inference tools attempt to highlight interactions of interest in various ways but do not attempt to improve the accuracy of their predictions beyond simply identifying them ((Dimitrov *et al*, 2021), Supplementary Table 1). However, scRNAseq data contains information about a cell type’s entire transcriptome, not only its expressed ligands and receptors. If one knew how each ligand affected its receiving cell’s gene expression profile, the receptor cell’s transcriptome could be used as evidence of ligand-receptor interaction and thus used to make more accurate cell-cell interaction predictions. We aimed to determine transcriptional signatures of ligand-receptor interaction to evaluate if they can be used generally to improve the accuracy of cell-cell interaction inference from scRNAseq data.

To determine the transcriptional signature of a ligand-activated receptor interaction, we used two complementary sources of data measuring transcriptional change in response to ligand perturbation. Connectivity Map is a database of over 1 million gene expression profiles in various cell lines in response to a variety of perturbands, including some secreted protein ligands (Subramanian *et al*, 2017). To efficiently measure this very large number of gene expression profiles, it uses a ligation-mediated amplification technique to measure the expression of 978 selected genes (referred to as “landmark” genes), from which the expression of a further 11,350 genes is inferred. To complement the breadth of this high-throughput resource, we also used published transcriptomics assays testing individual ligands perturbing a single cell type, originally curated by the authors of NicheNet (Browaeys *et al*, 2020). These individual microarray datasets were used to corroborate the conclusions drawn from the Connectivity Map data.

In investigating the transcriptional changes caused by each ligand treatment in the various cell lines available in these data sources, we observed that transcriptional responses to a ligand vary between cell types. Correlation of gene expression change upon treatment with a ligand was significantly worse comparing between cell types than within each cell type. Rarely were the genes whose expression was most responsive to ligand treatment in one cell type common across multiple cell types. This is not due to technical factors, as independent assays (biological replicates) within the same cell type often identify the same responsive genes. Finally, a non-linear machine learning method was not able to extrapolate predictions of ligand-treated versus untreated samples to those from a novel cell line. Our results suggest that additional cell-type-specific factors will be needed to improve the specificity of cell-cell interaction predictions from transcriptomic data, and highlights the importance of experimentally mapping cell-specific ligand-receptor interactions and their downstream transcriptional effects.

## Results

### Ligand-driven differential gene expression is not consistent across cell lines

To test if we could identify a characteristic gene expression signature downstream of a ligand-activated receptor, we used the Connectivity Map database of ligand-perturbed cell line gene expression profiles. Connectivity Map used the L1000 assay to measure the transcriptome of cell lines perturbed with individual ligands at given sets of concentrations and durations of treatment, reporting Z-scores for gene abundance comparisons to plate-matched controls (Subramanian *et al*, 2017). While the Connectivity Map uses the L1000 assay’s approximately 1000 (actually 978) measured gene abundances to infer gene expression values for most of the wider transcriptome, we used only the 978 measured Z-scores unless otherwise indicated. Connectivity Map’s 2017 release includes assays of 295 peptide ligands across 9 cell lines immortalized from a variety of tissues (Supplementary Figure 1). In these cell lines, ligands were often assayed in up to three biological replicates at a single concentration for a single duration of exposure. These data were used to assess the effect of cell context on the transcriptomic signature of ligand perturbation, unless otherwise indicated.

We first aimed to identify genes whose expression changed in response to ligand treatment consistently across all tested cell lines. To do this, differentially expressed genes were determined for each ligand perturbation by averaging Z-scores across all samples treated with the same ligand (Supplementary Figure 2). Statistical significance was determined by converting the mean Z-scores to p-values using the Gaussian distribution, and controlling the false discovery rate (FDR) by the Benjamini-Hochberg procedure. This resulted in the unexpected finding that most ligands did not affect gene expression in a sufficiently consistent manner across samples from multiple cell lines to result in statistically significant differential gene expression levels. Only 59 of the 295 assayed ligands caused significant changes in expression of any assayed genes at FDR ≤ 0.05, and only 19 ligands caused differential expression of more genes than expected by chance (p ≤ 0.05 by permutation testing, Figure 1). If one were to use these differentially expressed genes to build signatures of ligand-mediated transcriptional response, another issue becomes apparent - that relatively few genes are differentially expressed. Of the 978 genes whose expression was measured in these L1000 assays, 58 were significantly differentially expressed in response to any ligand, and only 32 of these were unique to a single ligand. These unique genes marked only 7 of the 295 assayed ligands, with Interleukin 19 (IL19) being notable for causing significant transcriptional changes in 23 genes not significantly affected by other ligands. Overall, it appears that most ligands in this dataset lack genes that are uniquely and significantly differentially expressed upon ligand treatment.

**Figure 1.**
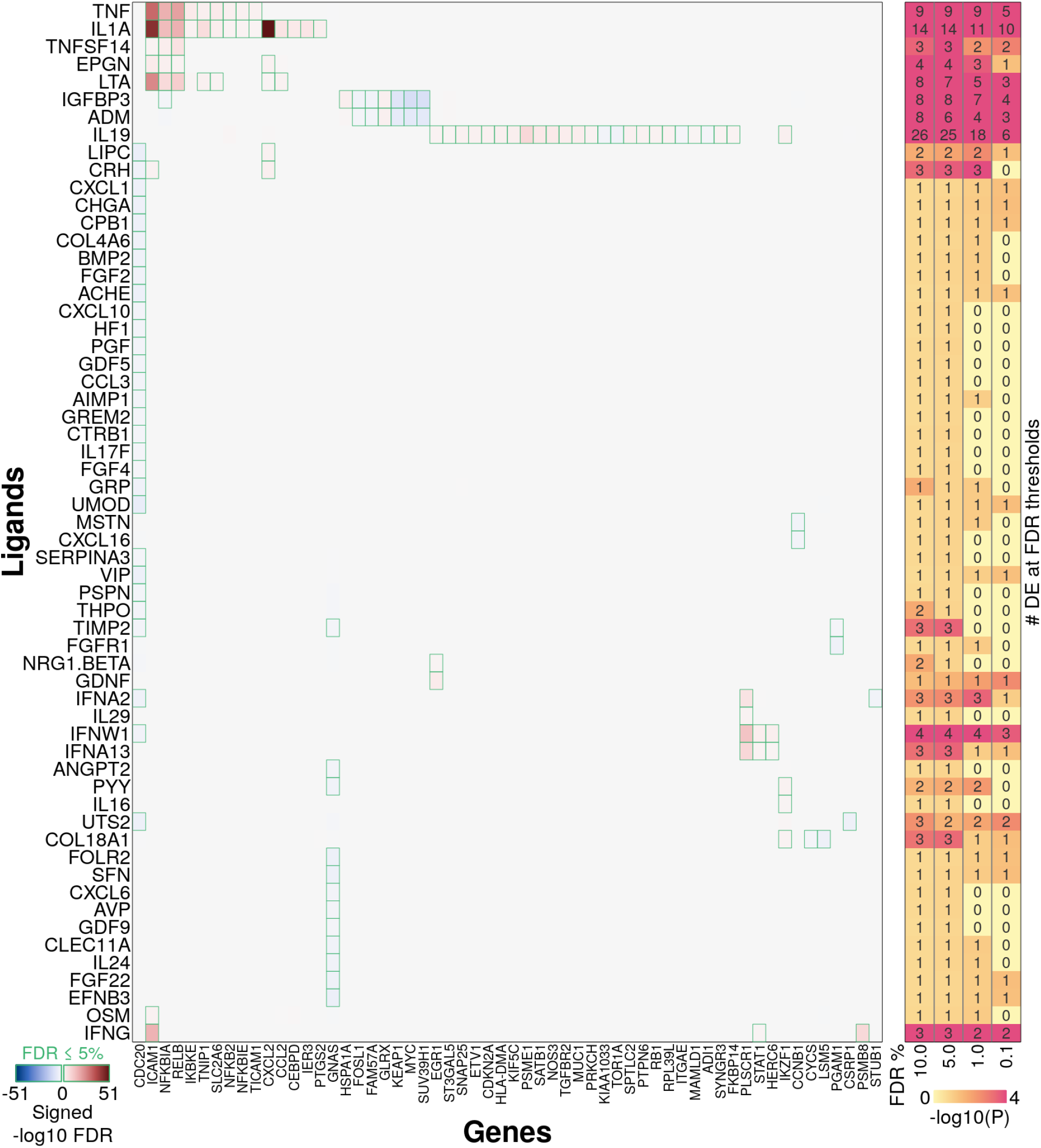
Few ligands drive consistent changes in gene expression across cell lines. A heat map showing average change in gene expression caused by ligand treatment averaged across all samples treated with the same ligand. Colours represent -log10 signed, false discovery rate corrected p-values derived from averaging Z-scores of gene expression change across all samples treated with the same ligand. Darker red represents a significant increase in gene expression, while darker blue represents a significant decrease in expression. All 59 (of 295) ligands where at least one gene was significantly different at an FDR threshold of 5% are shown. Green outline on a heat map cell represents a corrected p-value ≤ 0.05. Significantly differentially expressed genes unique to specific ligands could be used as transcriptional markers of ligand activity. However, the presence of vertical columns representing genes responsive to multiple ligands, and the overall sparsity of this heat map suggest that very few ligands possess such marker genes. On the right is a second heat map showing the probability of detecting the indicated number of differentially expressed genes by chance. The values indicate the number of significantly differentially expressed genes identified per ligand at the given FDR thresholds. Darker pink colour represents decreasing probability of seeing that number of differentially expressed genes by chance as calculated against a background distribution of differentially expressed gene counts determined by permuting sample labels.

### The lack of common ligand-responsive genes across cell contexts in the Connectivity Map dataset is not due to unexpected technical variability

To examine whether the Z-scoring procedure may have affected our result, we analyzed quantile-normalized gene abundance measurements (from which ligand-treatment Z-scores were calculated). These gene abundance measurements showed strong pairwise correlations between replicate experiments (mean Spearman correlation coefficient of 0.88, Supplementary Figure 3a), as did those from samples from the same cell line, whether or not they were treated with the same ligand (mean SCC of 0.85). Pairwise correlations between samples treated by the same ligand were notably less well correlated (mean SCC of 0.72) unless considering only samples from the same cell line (mean SCC of 0.86). Correlation between samples based on Z-scores was far worse (mean SCC of 0.08 between replicates, Supplementary Figure 3b), likely due to the relatively small proportion of the transcriptome responding to ligand treatment, as seen in the differential gene expression analysis above. Nevertheless, we observe the same overall pattern in analyses with both normalized gene abundance measurements and Z-scores, that pairwise correlation is improved when limiting the analysis to specific cellular contexts. Specifically, 213 of 295 ligands (72%) had significantly better (p ≤ 0.05 by Wilcoxon rank-sum test) mean pairwise correlations within at least one cell line versus across all lines, and 281 of 295 (95%) had significantly better correlations in at least one replicate set (same cell line, dosage, and duration of treatment) (Supplementary Figure 3c). Furthermore, there was a significant difference in the pairwise correlation of ligand-mediated transcriptional responses, as measured by Z-scores, across all samples treated with the same ligand, and those specifically from the same cell line, or same replicate set (Wilcoxon signed-rank p < 2.2e-16, Supplementary Figure 3d). Thus, considering cell context improves the correlation between changes in transcriptomes responding to the same ligand and this is not due solely to technical factors.

### Differential gene expression in response to treatment with an individual ligand is consistent within the same cell type context

After finding that correlation between ligand-treated transcriptomes improved when considering cell context, we used our differential gene expression analysis framework (Figure 1) to ask whether ligand mediated differentially expressed genes were consistent within the same cell context. We defined cell context by averaging gene expression Z-scores across ligand-treated samples from the same cell line across all experiments. In this analysis, in contrast to the analysis across all cell lines, all ligands cause statistically significant (FDR-corrected p-value ≤ 0.05) changes in the expression of at least one gene in at least one of the nine tested cell lines. This is consistent with the published Connectivity Map signatures (available at https://clue.io/), which are broken down by cell line. Nearly half (123 / 295) of the tested ligands caused differential expression (FDR-corrected p-value ≤ 0.05) of more genes than expected by chance in at least one cell line (p ≤ 0.05 by permutation testing), and 31 ligands caused a significant number DE genes in more than one cell line (Figure 2a). This is many more than the 19 ligands with more differentially expressed genes than expected by chance observed above (Figure 1). To quantify this context-dependent improvement in consistency of transcriptional response to ligand perturbation, we compared distributions of the probability of detecting as many differentially expressed genes by chance for each ligand when averaged across all samples versus the same, but for samples from the same cell line or biological replicate (Figure 2b). This comparison was made instead of comparing numbers of differentially expressed genes because when considering cell type context we are averaging across fewer samples, and thus would expect higher magnitude average Z-scores by chance. We avoid this bias by permuting sample labels to generate null distributions of Z-scores averaged across the appropriate number of samples, and calculating a p-value for the number of differentially expressed genes detected. When considering cell line context (i.e. averaging within the same cell line) these p-values improved for 274 (93%) of the 295 ligands (Supplementary Figure 4), and overall it was significantly less likely that the number of differentially expressed genes per ligand was detected by chance (p < 2.2e-16 by Wilcoxon rank-sum test). This finding was robust to the FDR threshold used to determine the number of differentially expressed genes. Similar results were obtained when including the duration and dosage of ligand treatment along with cell line context - i.e. averaging Z-scores within replicates (p < 2.2e-16 by Wilcoxon rank-sum test compared to per ligand, no change compared to per ligand in the same cell line, Figure 2b). Thus, for most ligands tested in Connectivity Map, gene expression changes are dependent on cell context.

**Figure 2.**
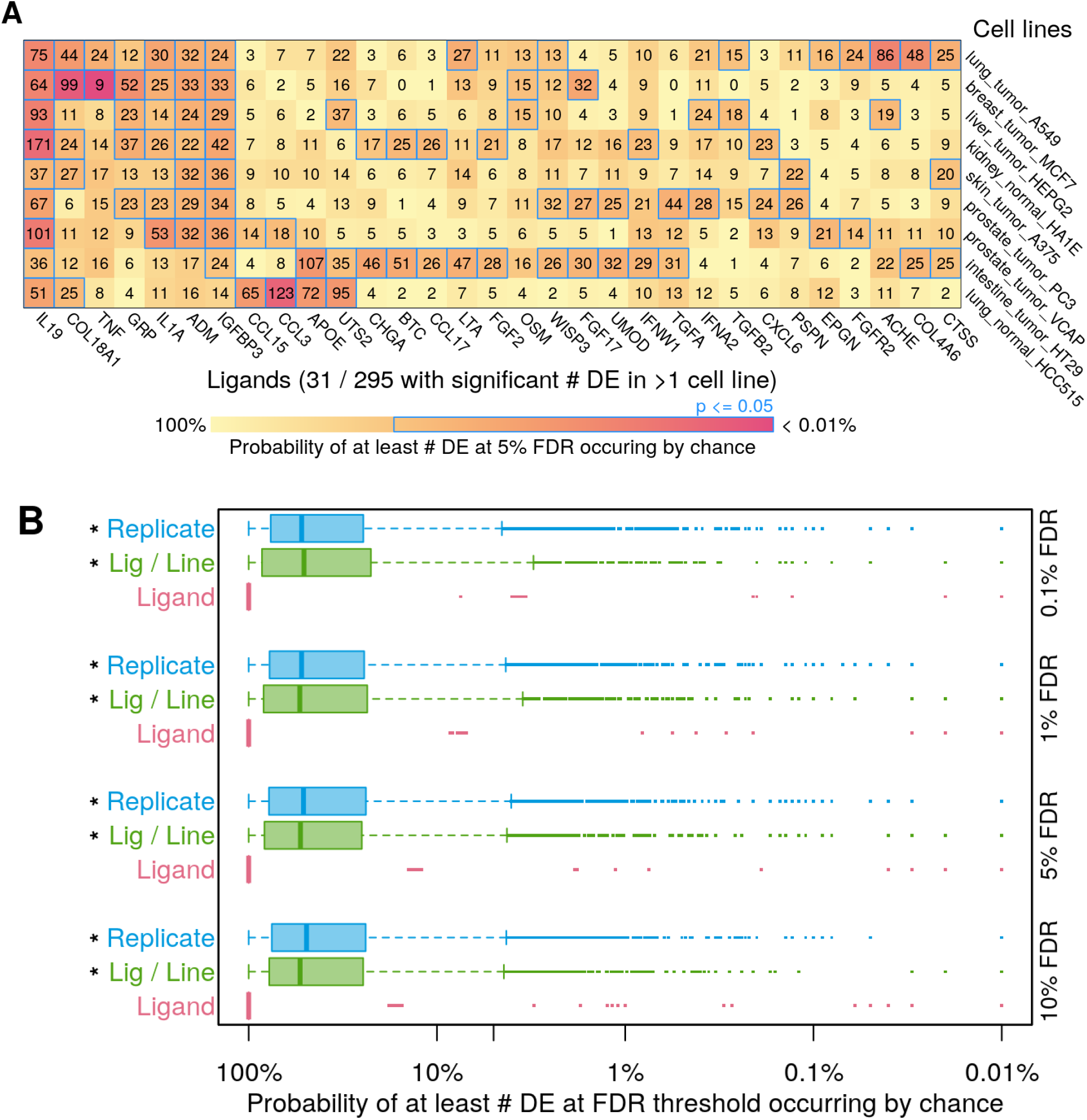
Many ligands drive consistent changes in gene expression in some individual cell lines. **a)** A heat map showing significance scores for the number of differentially expressed genes (at FDR of 5%) caused by a ligand’s perturbation of each cell line. The count of differentially expressed genes in response to each ligand (column) in each cell line (row) is shown. Statistical significance (indicated by darker colour) is calculated by determining the probability of seeing that number of differentially expressed genes by chance against a background distribution of differentially expressed gene counts determined by permuting sample labels. Ligands are included in the heat map if they cause differential expression of a significant number of genes in more than one cell line (p ≤ 0.05 by permutation test). **b)** Boxplots showing the distribution of significance scores (as calculated in panel a) when averaging gene Z-scores across all samples treated with the same ligand (as in Figure 1), all samples treated with the same ligand in the same cell line (“Lig / Line”, from panel a), and all replicate samples (same ligand, cell line, dosage, and duration of treatment). Significance is represented as the probability of seeing as many differentially expressed genes by chance against a background distribution of differentially expressed gene counts determined by permuting sample labels. Numbers of differentially expressed genes were calculated at the False Discovery Rate (FDR) thresholds indicated on the right vertical axis, and * indicates that the center of that distribution is significantly different (p < 2.2e-16 by Wilcoxon rank-sum test) than when considering all samples treated with the same ligand.

### Modeling non-linear response to ligand stimulus does not improve the transcriptional signature generalizability

Using significantly differentially expressed genes, it is difficult to identify generalized (i.e. context-independent) transcriptomic signatures for most ligand-receptor interactions. It is possible that there are generalizable non-linear combinations of gene transcription responses to ligand perturbation that could also include genes that vary, but are not significantly differentially expressed in the context of whole transcriptome statistical tests. Random forest models can make use of non-linear information, as they use votes from a series of decision trees to build classifiers. They have also proven to be useful as tools for feature selection and classification tasks in transcriptomics, and achieve state of the art performance in these tasks (Xu *et al*, 2020; Acharjee *et al*, 2020; Kursa, 2014). To assess the ability of a random forest model to identify ligands from their transcriptomic profile, models were trained for each ligand across all assayed cell lines for the binary classification task of determining whether the sample was treated with the given ligand. A subset of 16 ligands (BTC, EGF, FGF1, GAS6, GDNF, HBEGF, HGF, IFNG, IGF1, IGF2, IL17A, IL4, IL6, INS, TGFA, and TNF) with higher sample numbers were used for this task to support good model power. These ligands were tested in the 9 cell lines used in the Connectivity Map data above and assayed at multiple concentrations and exposure durations in an additional 5 breast cancer cell lines. The models’ performance on this task was generally poor (Figure 3a - rows with CTRL prefix), except when predicting IFNG and TNF treatment, which agrees with our differential gene expression results. To determine whether the models’ prediction ability is generalizable across cell types, or conversely whether transcriptional responses to ligands are cell-type specific, these models were trained again while withholding data from one cell line, and performance was tested on the withheld cell line (Figure 3a). Performance decreased when extrapolating to a novel cell line, except in the case of IFNG (Figure 3b, pairwise AUPR decreased significantly (p < 1e-16) by Wilcoxon signed-rank test). This is consistent with our previous finding that both correlation in transcriptional change and number of significantly differentially expressed genes increases more than expected by chance when considering cell-type specific responses to a ligand stimulus, compared to considering a generalized response across cell types.

**Figure 3.**
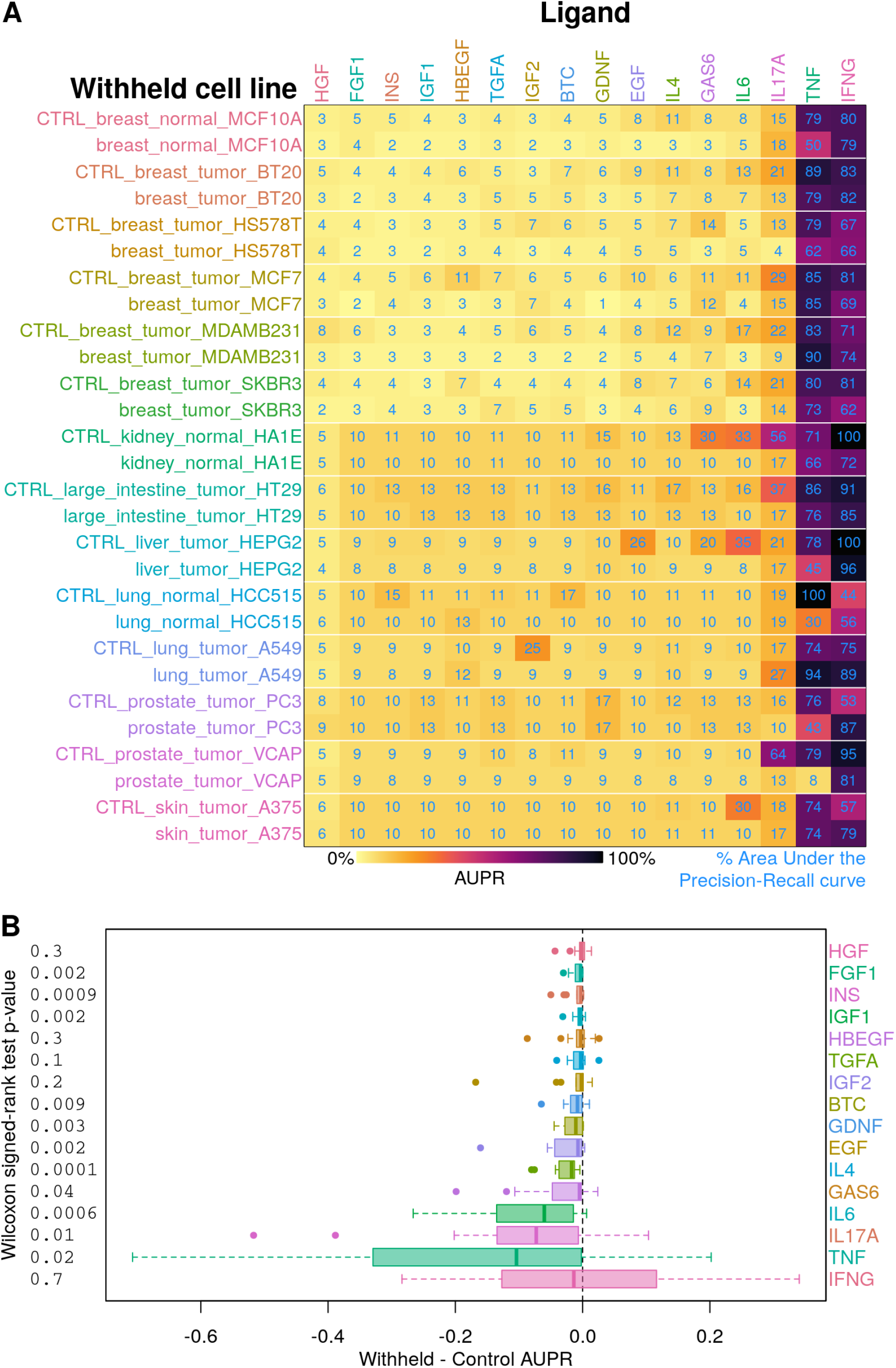
Extrapolating to novel cell lines reduced performance of machine learning models trained to identify ligand-treated transcriptomes. **a)** Each column displays the results of a series of random forest models trained to identify transcriptomes treated by the indicated ligand, and each row indicates the performance of the given model in the cell line that was withheld for training and used for testing, or the respective control (“CTRL”), where equal numbers of samples from all cell lines were used for training and testing. Random forest model performance represented by Area Under the Precision-Recall curve (AUPR, darker colour is higher performance, percentage values shown in blue text). Columns are ordered by increasing mean control AUPR. **b)** Boxplots showing change in AUPR between training on all cell lines and training on all but a withheld cell line, and predicting on the withheld cell line. Boxplots are ordered as columns in the above heat map by increasing mean control AUPR to give context to the possible magnitude of change in AUPR. Data to the left of the dashed vertical line indicate that holding out a cell line reduces model performance, while points to the right of the dashed line indicate improvement. Wilcoxon signed-rank test was used to determine significance of pairwise change in accuracy, p-values reported to the left of each boxplot.

A complementary classification task is to identify which ligand treatment drives each set of transcriptional changes. A different set of random forest models were trained to perform this task and this generated similar results to the previous random forest analysis (Supplementary Figure 5). Median accuracy when extrapolating to novel cell lines was reduced compared to the already poor accuracy (32%) when training on all cell lines (5% decrease in median accuracy, p = 1.3e-4 by Wilcoxon signed-rank test).

One possible explanation for the general difficulty in establishing transcriptional signatures of ligand response (either linear or nonlinear) is poor signal in this data. This may indicate a lack of response to the ligand perturbation, possibly because the ligand’s cognate receptor is not expressed in that cell line. To assess this possibility, receptor expression was correlated with ligand prediction accuracy in our random forest model tasks (Supplementary Figure 6). Cognate receptors for each ligand were determined from a database of known ligand-receptor interactions (Ximerakis *et al*, 2019), and their quantile-normalized gene abundance per cell line were calculated from the Connectivity Map control data. While epidermal growth factor (EGF) receptor family members showed reasonable correlation with prediction accuracy of transcriptional response to the EGF ligand family member TGFA, for most ligands the correlation between cognate receptor expression and ligand prediction accuracy was unclear. Thus, we do not find that receptor expression level explains ligand perturbation signal in general.

### Cell-type specific transcriptional response to ligand perturbation is corroborated by independent microarray assays

To corroborate our main finding from the Connectivity Map database that transcriptional response to ligand perturbation is cell-type dependent, we repeated our analysis on a complementary transcriptomic dataset. The authors of the cell-cell interaction prediction method NicheNet (Browaeys *et al*, 2020) used the NCBI Gene Expression Omnibus (Barrett *et al*, 2013) to curate a collection of microarray experiments in which transcriptional response to ligand stimulation was assayed. Twenty ligands appeared more than once in the collected data, of which eleven ligands were assayed in more than one independent experiment from different cell types and thirteen were assayed more than once in the same cell type by independent labs in separate experiments (Supplementary Figure 7). As was done with the Connectivity Map data, we calculated pairwise Spearman correlation coefficients between all fold-change values from experiments where the same ligand was used to perturb cells (Figure 4a). As with the Connectivity Map data, change in gene expression upon ligand treatment in different cell lines was poorly correlated (mean Spearman correlation coefficient of 0.1). Ligand-induced gene expression changes were significantly better correlated when comparing independent experiments in the same cell type (mean Spearman correlation coefficient of 0.35, p = 3.4e-7 by Wilcoxon rank-sum test). This mirrors our findings analyzing the Connectivity Map data, albeit with a much stronger effect and better coverage of the whole transcriptome. By comparing overall ligand-induced changes in gene expression within and between cell types, the published transcriptomic data clearly support the finding that transcriptional response to ligand treatment is highly dependent on cell type.

**Figure 4.**
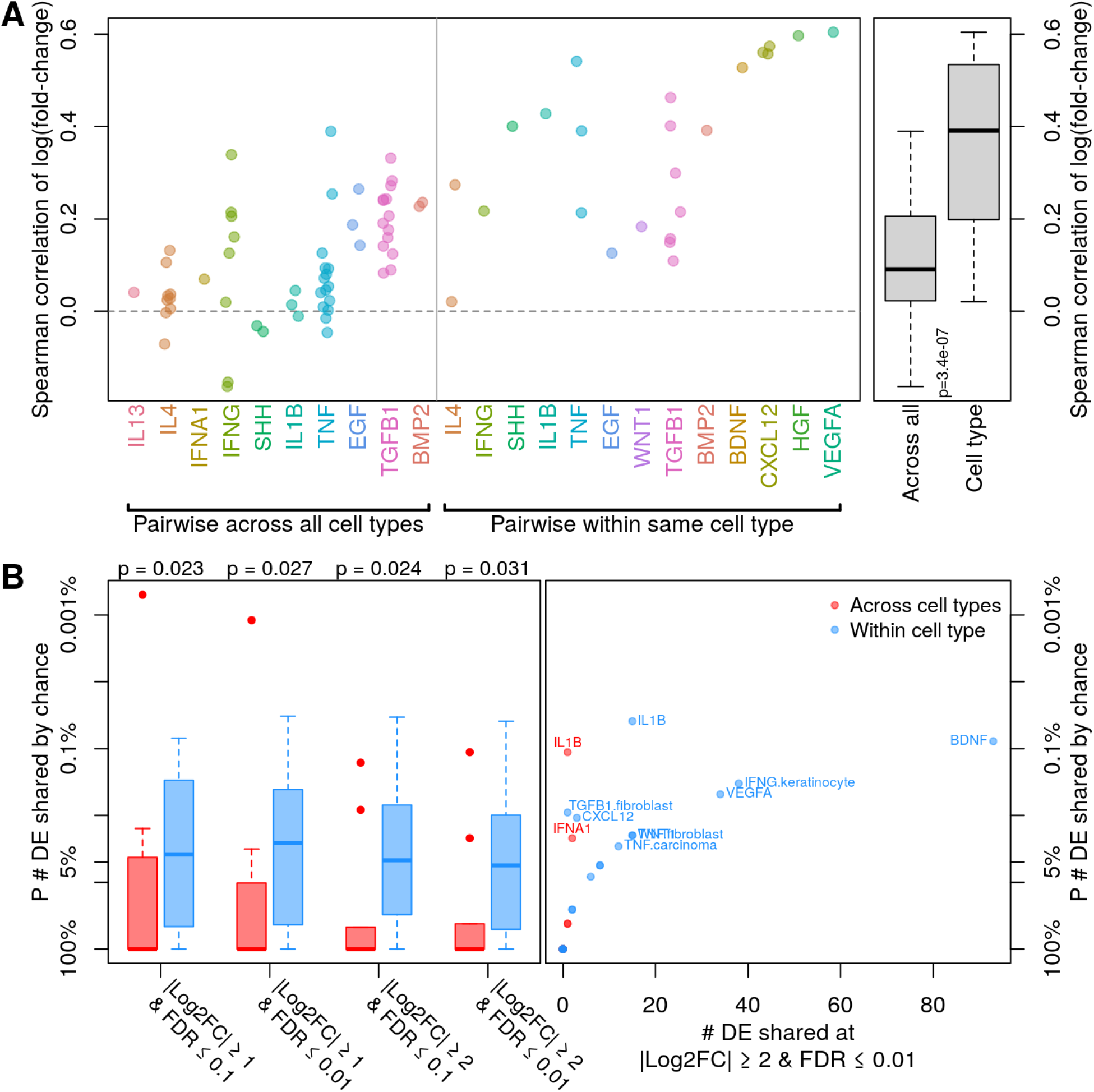
Context-specific response to ligand stimulus is corroborated by independent transcriptomics assays. **a)** A strip plot showing pairwise Spearman correlations between log fold-change values from NicheNet-curated transcriptomics assays assessing change in gene expression caused by ligand treatment, for all pairs of assays for each ligand (left) or pairs of assays using the same cell type (right). Correlations in gene expression change between samples in different cell types treated with the same ligand are shown on the left, with correlations between samples from the same cell type and ligand treatment on the right. The data is summarized in the boxplot on the far right, which indicates that ligand-dependent transcriptional changes are significantly more correlated in the same cell type (p = 3.4e-7 by Wilcoxon rank-sum test). **b)** Ligand-dependent gene expression changes from published transcriptomics assays common across all assays. On the left are box plots showing the statistical significance of overlapping differentially expressed genes (y-axis), using various cutoffs to define differential expression (x-axis). Statistical significance is represented as the probability of seeing such overlap occurring by chance, as calculated by permutation testing. Indicated p-values (top) are from Wilcoxon rank-sum tests comparing probabilities of overlap by chance between samples from different cell types (red) versus within the same cell type (blue). On the right is a representative scatter plot showing the data from one pair of boxplots, with the x-axis representing the number of differentially expressed (DE) genes shared between all datasets treated with the same ligand (in red) or all datasets from the same cell type and treated with the same ligand (in blue). Ligands or ligand and cell type were labeled if the overlap p-value was ≤ 0.05.

We also used the NicheNet-curated gene expression data to assess the finding that some genes display significant and consistent changes in gene expression upon ligand treatment, but these are context-specific. Significantly differentially expressed genes in response to ligand treatment were defined from each ligand-perturbed microarray dataset at multiple effect size and statistical significance thresholds to ensure robustness of findings. The number of significantly differentially expressed genes shared across all datasets treated with the same ligand, or treated with the same ligand in the same cell type, was then counted, and the probability of those numbers of shared DE genes occurring by chance was calculated by permutation testing (Figure 4b). The chance of differential gene overlap between datasets was significantly less likely when considering ligand-induced changes in gene expression within each cell type, irrespective of the fold change and false discovery rate thresholds used to define significant differential expression (Figure 4b right panel, p < 0.05 by Wilcoxon rank-sum tests). This is consistent with the finding from Connectivity Map data analysis that the vast majority of ligands show cell line specificity in transcriptional response (Figure 2b).

From this analysis, we find that only two ligands, IL1B and IFNA1, share a significant number of ligand-responsive differentially expressed genes across all cell types in the NicheNet-curated data. None of these genes were differentially expressed across cell types in the Connectivity Map data(Supplementary Figure 8). Of the ligands assayed in NicheNet-curated transcriptomics data, TNF and IFNG caused the most generalizable transcriptional changes in the Connectivity Map analysis (respectively 9 and 3 significantly differentially expressed genes at 5% FDR, Figure 1). The most consistently differentially expressed gene in response to both TNF and IFNG is ICAM1, but it is not upregulated in all the TNF-treated cell lines, lacking significant change in expression in both smooth muscle and immune cell contexts from the NicheNet-curated data, and in the prostate tumour line VCAP from Connectivity Map (Figure 5). Two other consistently upregulated genes in Connectivity Map in response to TNF, NFKBIA and RELB, were similarly inconsistent, with no significant differential expression in the breast cell line MCF10A or prostate tumour line VCAP in Connectivity Map. In response to IFNG, ICAM1 was upregulated in many, but not all, cell contexts. It showed no significant response in immune or fibroblast cells from the NicheNet curated data, nor in the breast tumour line HS578T from Connectivity Map (Supplementary Figure 9). Thus, even the most consistent transcriptional response signatures across many independent data are not always observed, emphasizing that cellular context is still important even in these special cases.

**Figure 5.**
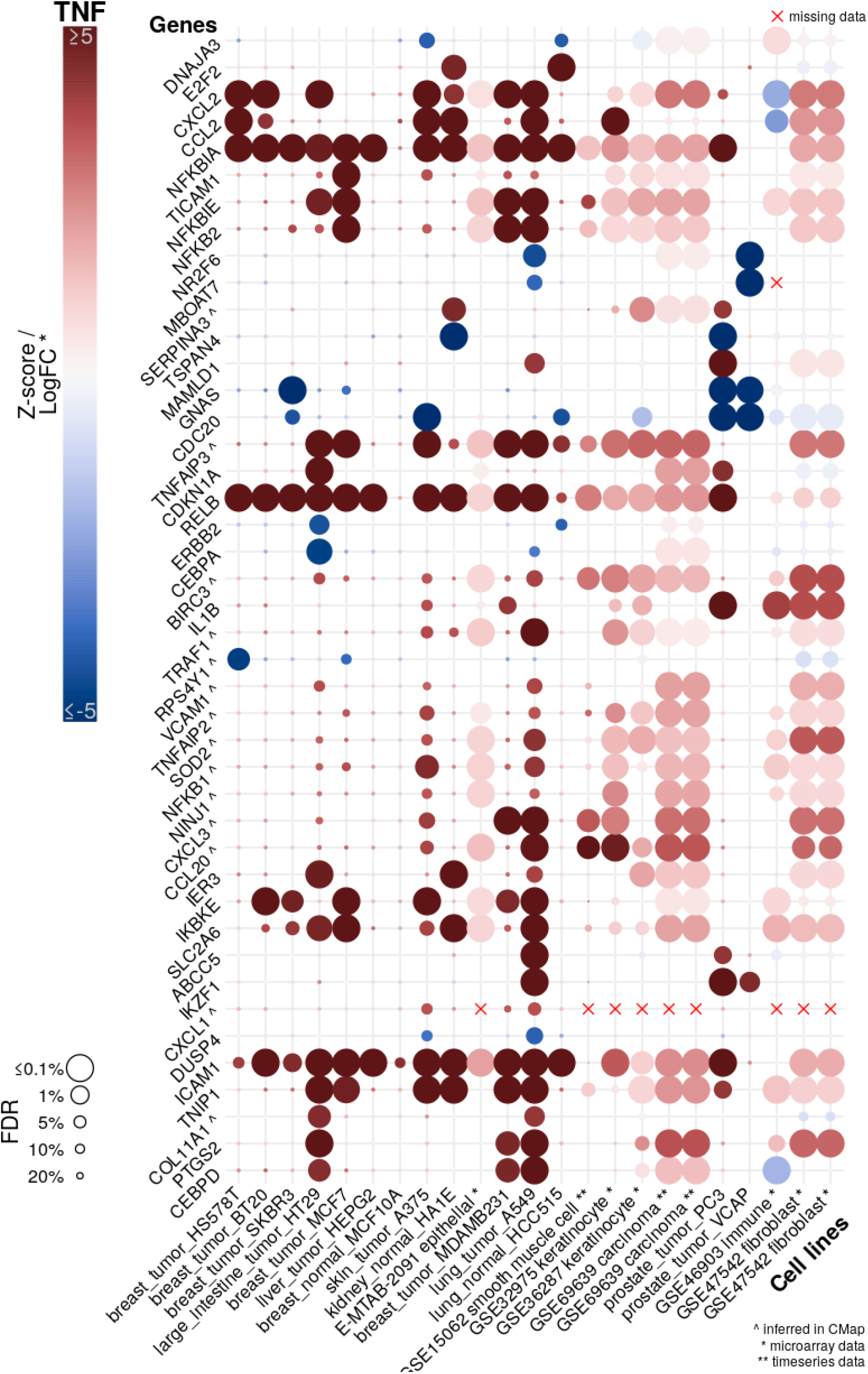
Gene expression changes in response to TNF ligand stimulus differ between cell types. A matrix plot showing mean change in gene expression per cell line upon TNF treatment for selected genes from both Connectivity Map and NicheNet-curated transcriptomics data. Cell lines are on each column, with NicheNet-curated transcriptomes indicated with *, and those from time series experiments where only magnitude of gene expression change was recorded with a **. Genes (in rows) were included if they were significantly differentially expressed (at 10% FDR) in at least two Connectivity Map cell line assays or the maximum possible number of overlapping NicheNet-curated cell line assays. Genes whose expression was inferred in Connectivity Map are indicated with ^. Genes whose expression was not reported in a given experiment are indicated with a red x instead of a dot. Dot colour indicates difference in gene expression upon ligand treatment, measured as Connectivity Map Z-score and log2 fold-change from NicheNet-curated transcriptomes. Dot size indicates false discovery rate-corrected significance. If a gene were consistently differentially expressed in response to ligand treatment, an unbroken horizontal line of circles of the same colour would appear on this plot.

## Discussion

If the various cell types of a tissue analyzed by single-cell RNAseq are communicating through ligand-receptor interactions, one may expect to see evidence of these ligand-mediated signaling events in the receiving cell’s transcriptome. In this study, two independent and complementary transcriptome assay databases measuring gene expression change in response to ligand perturbation were used to investigate this hypothesis. Within each cell type tested, change in gene expression in response to each ligand was often consistent and predictable, as evidenced by nearly half of the ligands in the Connectivity Map data causing significant differential expression of more genes than expected by chance. However, ligand response across cell types was generally inconsistent and unpredictable. These results have important implications for cell-cell interaction prediction from scRNAseq data, which cannot rely on generalizable, characteristic and easily predictable ligand response signatures to identify receptors that are actively signaling to effect downstream transcriptional changes. Instead, only cell-type-specific ligand response signatures will be useful. Unfortunately, this cellular context dependent ligand response information is not available for most cell types.

We also found that certain genes are activated by many tested ligands, as evidenced by the vertical lines of significant differential expression in Figure 1. That independent experimental perturbations can have similar transcriptional effects has been noted before, most notably by Crow *et al*., who built a prior probability of differential expression for all genes with 80% accuracy across a wide variety of transcriptomic datasets (Crow *et al*, 2019). While one might attempt to mitigate the apparent redundancy in transcriptional response to ligand perturbation by making more coarse-grained predictions by ligand family, Crow *et al*. suggests that by virtue of either the way transcriptomic data is collected, or biological redundancy in transcriptional response to stimuli, that may not be sufficient to ensure uniqueness of ligand signaling signatures.

Cell-cell communication inference is a popular method for understanding tissue function from scRNAseq data, but no current method is able to address the challenge we describe here (Supplementary Table 1). It may be possible to infer cell-type-specific signaling networks downstream of a given receptor using transcriptome data, but this is currently an unsolved problem (Larsen *et al*, 2019). The closest any cell-cell prediction method comes to addressing this problem is described by Browaeys *et al*. in their NicheNet method, which uses a combination of published signaling and gene regulatory pathways within a cell to predict the expected transcriptional response to ligand perturbation (Browaeys *et al*, 2020). Unfortunately, current pathway databases do not consider cell type context in their networks, and while the literature acknowledges that gene regulatory networks are cell type specific, one needs context-specific training data to predict gene regulatory networks with such specificity (Chasman & Roy, 2017), (Margolin *et al*, 2006). As such, NicheNet lacks the prior knowledge resources to account for the context-dependent nature of transcriptional response to ligand perturbation. Furthermore, testing these predictions requires ground-truth data representing the cell types of interest, and the efforts of the NicheNet authors have highlighted the paucity of such data (Browaeys *et al*, 2020).

Ultimately, current cell-cell interaction predictions can provide a non-specific list of potential ligand-receptor pairs as a hypothesis generation exercise, but study authors must test these predictions to both identify true sources of cell-cell signaling, and provide the sort of ground-truth data necessary for further improvement of these predictions.

## Materials & Methods

### Data and code availability

All analysis was performed in the R statistical programming language (R Core Team, 2018), with all code available at https://github.com/BaderLab/Brendan_CCInxPred. Connectivity Map data is deposited in NCBI GEO with accession GSE92742 (Subramanian *et al*, 2017). NicheNet-curated data is from the ‘expression_settings.rds’ file deposited in Zenodo as record 3260758 (Browaeys *et al*, 2019).

### Connectivity Map differentially expressed gene expression analysis

Connectivity Map Z-scores for changes in gene expression upon ligand treatment were averaged across all samples treated with the same ligand, same ligand in the same cell line, or from the same technical replicate set. False Discovery Rate (FDR)-corrected p-values were calculated from averaged Z-scores using the Gaussian distribution. A background distribution of FDR-corrected p-values was generated by averaging each gene’s Z-scores across a random set of samples of equivalent size. This background distribution was used to calculate the probability of seeing as many differentially expressed genes by chance for each set of samples. The R scripts to perform these calculations and generate Figure 1 and Figure 2 are ‘lig295_DE_FDR.R’ and ‘DEoverlap_FDR_forPaper.Rmd’ in the GitHub repository cited above.

### Connectivity Map sample correlations

Spearman correlation coefficients were calculated for all pairs of samples from the same cell line, treated with the same ligand, same ligand in the same cell line, or from the same technical replicate set. Differences in the distributions of pairwise correlation coefficients were tested using the Wilcoxon rank-sum test. The R scripts to perform these calculations and generate Supplementary Figure 3 are ‘Zcorr.R’ and ‘Zcorr_forPaper.Rmd’ in the GitHub repository cited above.

### Connectivity Map random forest modelling

Random forest models were trained for each ligand using *ranger* to classify samples as either treated with the ligand or not (Wright & Ziegler, 2017). Training sets were balanced by randomly selecting an equivalent number of samples treated with other ligands to represent the negative case for each ligand. “Control” models were trained on samples from all cell lines, while “test” models for each cell line were trained on all lines except the cell line in question. The models were tested on all withheld data. The R scripts to perform these calculations are generate Figure 3 are ‘200813_newVall_probs_lvl4.R’ and ‘RFresults_forPaper.Rmd’ in the GitHub repository cited above.

Random forest models were also trained on the same datasets as above to identify which of the 16 ligands each sample was treated with. The R scripts to perform these calculations and generate Supplementary Figure 5 are ‘200524_lvl5_mixall_leave1out.R’ and ‘LigPred_extrapolate.Rmd’ in the GitHub repository cited above.

### Connectivity Map receptor availability analysis

Average of quantile-normalized gene expression values of cognate receptors for each ligand in each cell line were correlated with accuracy of binary classification by the relevant random forest model. Cognate receptors for each ligand were identified from the ligand-receptor database used by *CCInx* (Ximerakis *et al*, 2019). The R script to perform these calculations and generate Supplementary Figure 6 is ‘RecExpr.Rmd’ in the GitHub repository cited above.

### NicheNet database correlation and differential expression analysis

Spearman correlation coefficients between log-scaled fold-change values of gene expression difference between treated and untreated samples were calculated for all pairs of samples treated with the same ligand, or treated with the same ligand in the same cell type. Differences in the distributions of pairwise correlation coefficients were tested using the Wilcoxon rank-sum test.

Differentially expressed genes in each sample were determined using multiple fold-change and FDR thresholds. To determine the statistical significance of overlap between differentially expressed gene lists from samples treated with the same ligand, or treated with the same ligand in the same cell type, the null probability of overlap was calculated by permuting sample labels. Differences in the distributions of overlap probabilities were tested using the Wilcoxon rank-sum test.

The R scripts to perform these calculations and generate Figure 4 are ‘NicheNet_calc.R’ and ‘NicheNet_forPaper.Rmd’ in the GitHub repository cited above. The R script to generate Figure 5, Supplementary Figure 8, and Supplementary Figure 9 is ‘NicheNet.Rmd’ in the GitHub repository cited above.

