## Supplementary figures & table for "Transcriptional signatures of cell-cell interactions are dependent on cellular context"

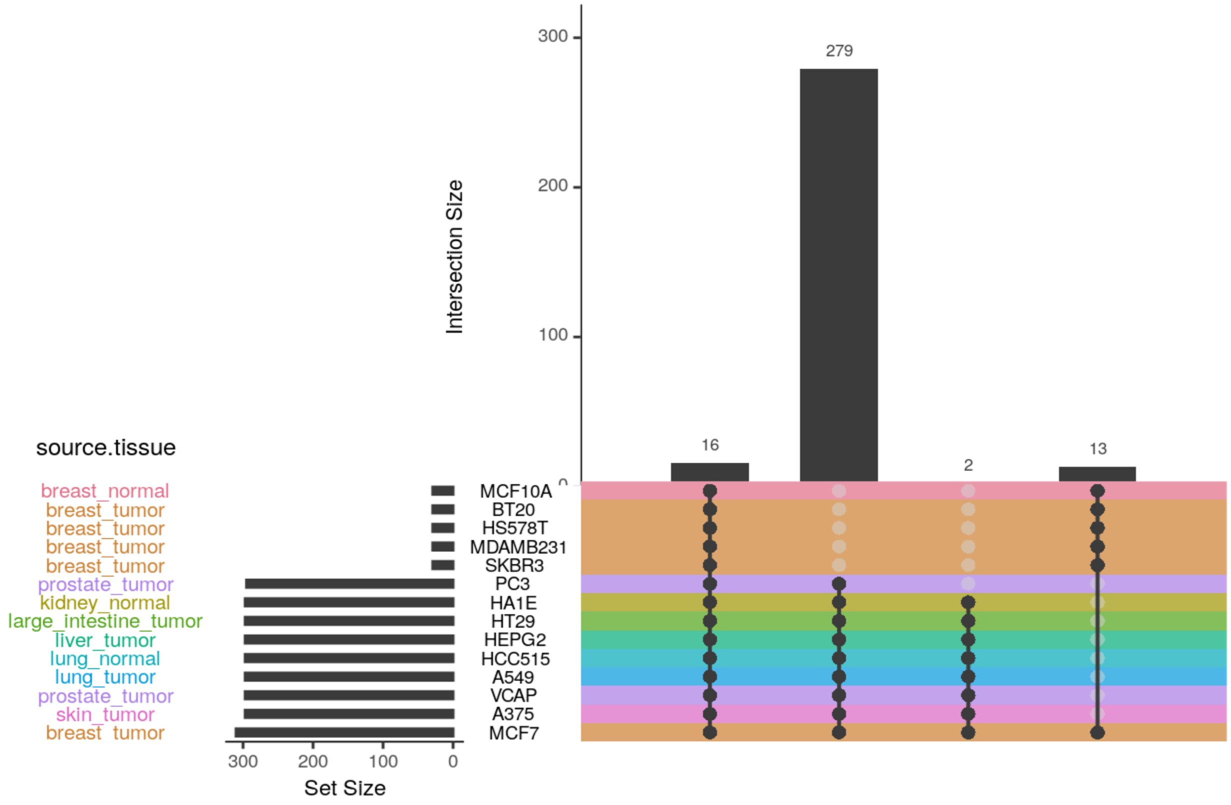


**Supplementary Figure 1. Summary of Connectivity Map ligand and cell line coverage.**

An UpSet plot (Lex *et al*, 2014) showing the distribution of ligands assayed in each of the 14 cell lines. 16 ligands were assayed in all 14 cell lines, and 295 ligands were assayed in 9 of the 14 cell lines, covering all seven tissue sites represented by the 14 cell lines.

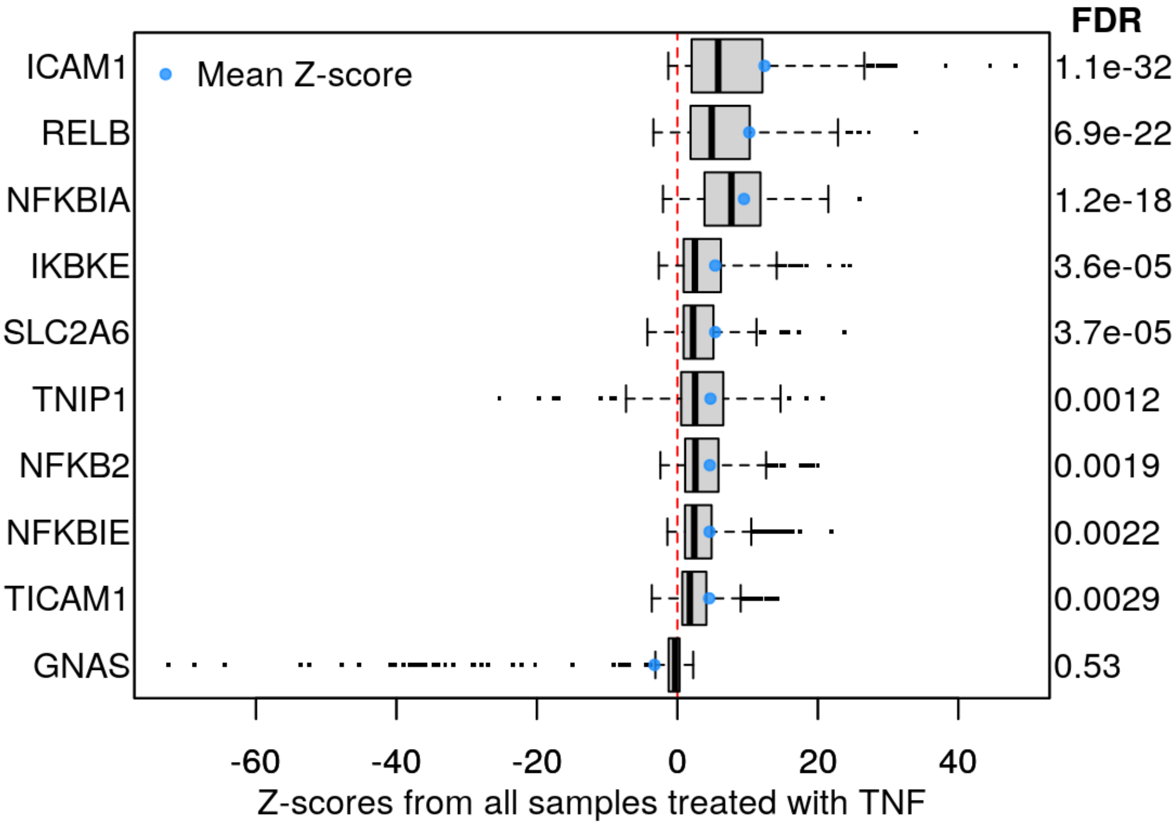


**Supplementary Figure 2. Average changes in gene expression upon TNF ligand treatment (Z-scores) across all samples in Connectivity Map.**

Shown here are the genes with the highest magnitude mean Z-scores (blue dot) for samples treated with TNF. To determine which gene's expression level was consistently perturbed across samples treated with the same ligand, Z-scores representing normalized change in gene expression for each treated sample were averaged. P-values for each change in gene expression were calculated from the mean Z-score then corrected for multiple testing by the Benjamini-Hochberg procedure to control False Discovery Rate (FDR).

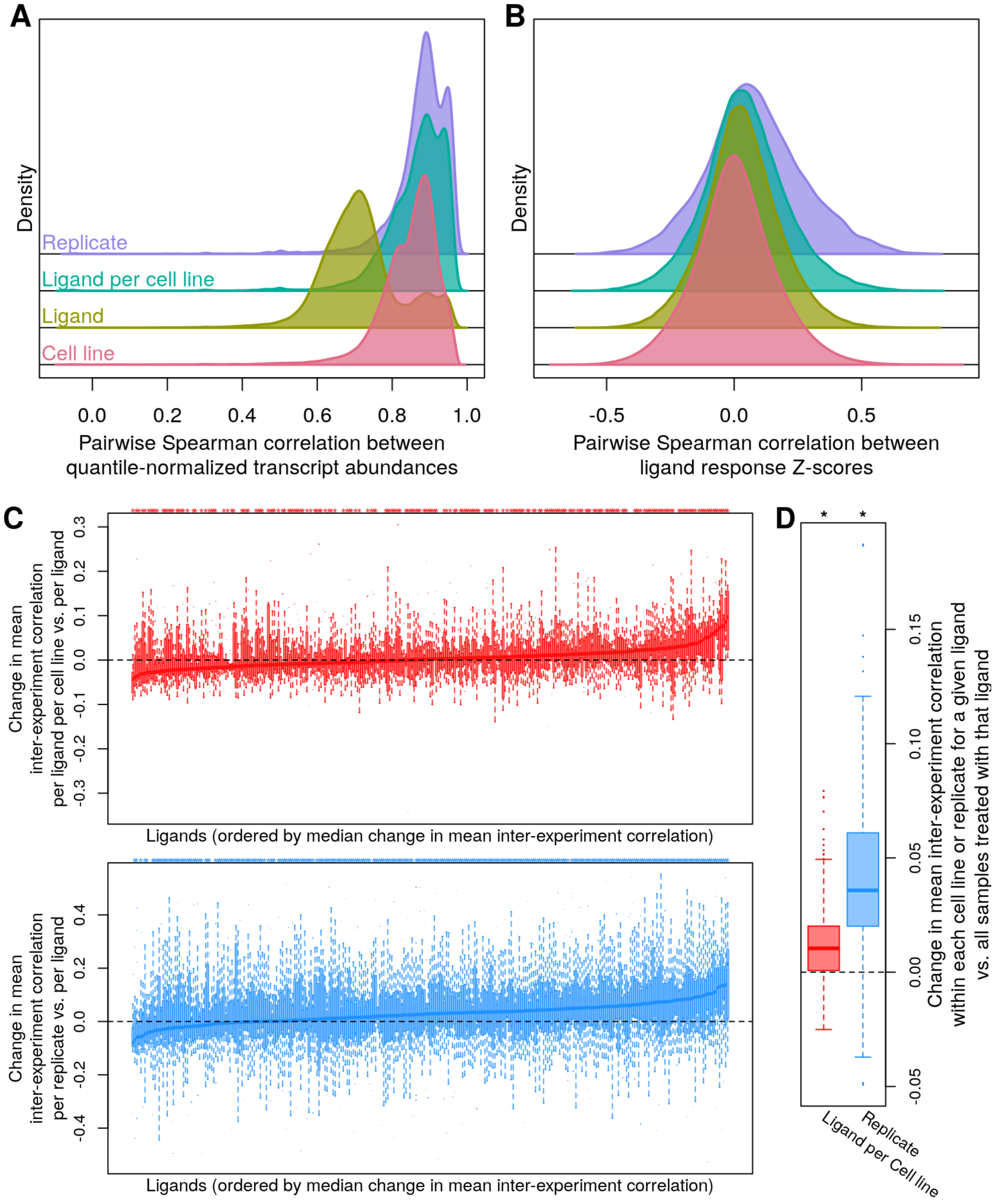


**Supplementary Figure 3. Correlation of change in gene expression upon ligand perturbation is decreased when not accounting for cell context, despite good correlation of gene expression between replicates.**

**a)** Distribution of pairwise Spearman correlation coefficients between quantile-normalized gene expression magnitude values for all sample pairs from the same cell line (red), treated with the same ligand (yellow-green), treated with the same ligand in the same cell line (teal), or from the same replicate set (same ligand, dosage and duration of treatment, in the same cell line; purple).
**b)** Distribution of pairwise Spearman correlation coefficients between Z-scores representing change in gene expression upon ligand treatment, for the same sets of samples as panel a.
**c)** Boxplots showing the change in mean pairwise Spearman correlation coefficients between Z-scores for each ligand when considering only correlations within the same cell line (top, red) or within the same replicate set (same cell line, dosage and duration of treatment ; bottom, blue). Each box represents a ligand, and the y-axis values represent the difference between the mean of all pairwise Spearman correlation coefficients between all samples treated with that ligand; subtracted from the means of all pairwise correlations between samples treated with that ligand in each cell line (red) or in each replicate set (blue). Ligands are ordered on the x-axis by the median value of these differences in means. The small * above the boxplots indicates that at least one cell line (red; 213 / 295 (72%) of ligands) or replicate (blue; 281 / 295 (95%) of ligands) had a significant improvement in mean inter-experiment Z-score correlation compared to the mean of all inter-experiment correlations for that ligand ($p \leq 0.05$ by Wilcoxon rank-sum test).
**d)** Boxplots summarizing the results in panel c. Each boxplot contains a data point for each of the 295 ligands. Each point is the difference between the mean of all inter-experiment correlations for a given ligand, subtracted from the mean of all inter-experiment correlations from each cell line (red) or each replicate set (blue). Correlations per ligand within each cell line, and per replicate are both significantly (* = $p < 2.2e-16$ by Wilcoxon signed-rank test) greater than across all samples treated with the same ligand.

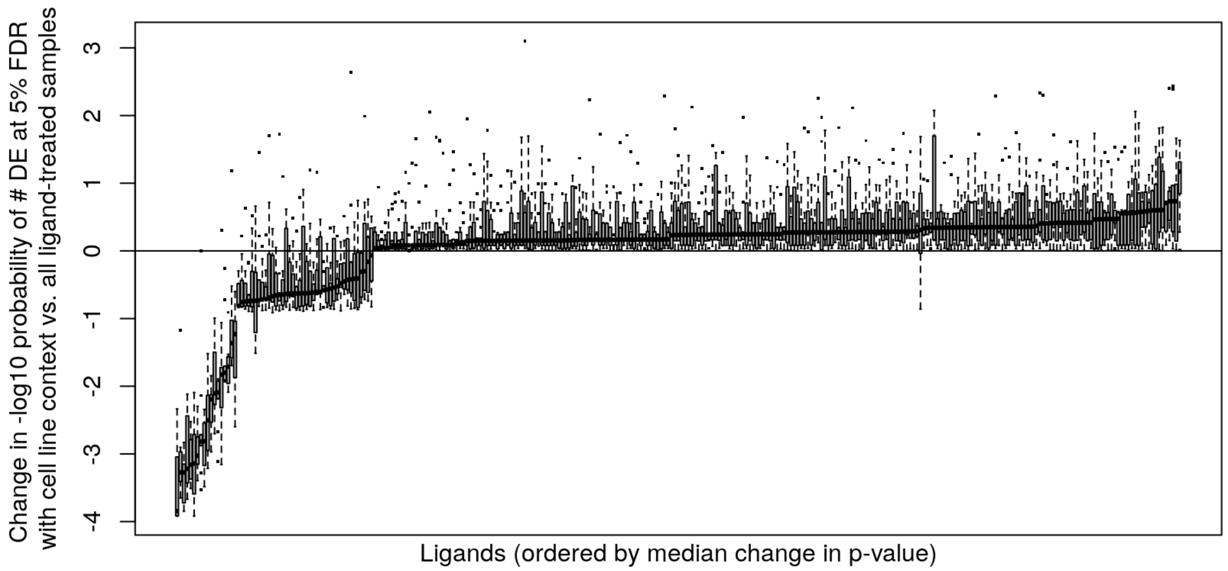


**Supplementary Figure 4. Significantly more differentially expressed genes are detected when considering cell line context for most ligands.** Each boxplot represents the change in significance score between averaging across all Z-scores from samples treated with a given ligand, subtracted from the averages of Z-scores from all samples treated with a given ligand in each cell line. Significance score is the negative log10 transformation of the probability of seeing that number of differentially expressed genes by chance, calculated using a background distribution of differentially expressed gene counts determined by permuting sample labels. Values above zero indicate that significantly more differentially expressed genes were caused by the given ligand in that cell line than that ligand causes to be differentially expressed generally across all cell lines.

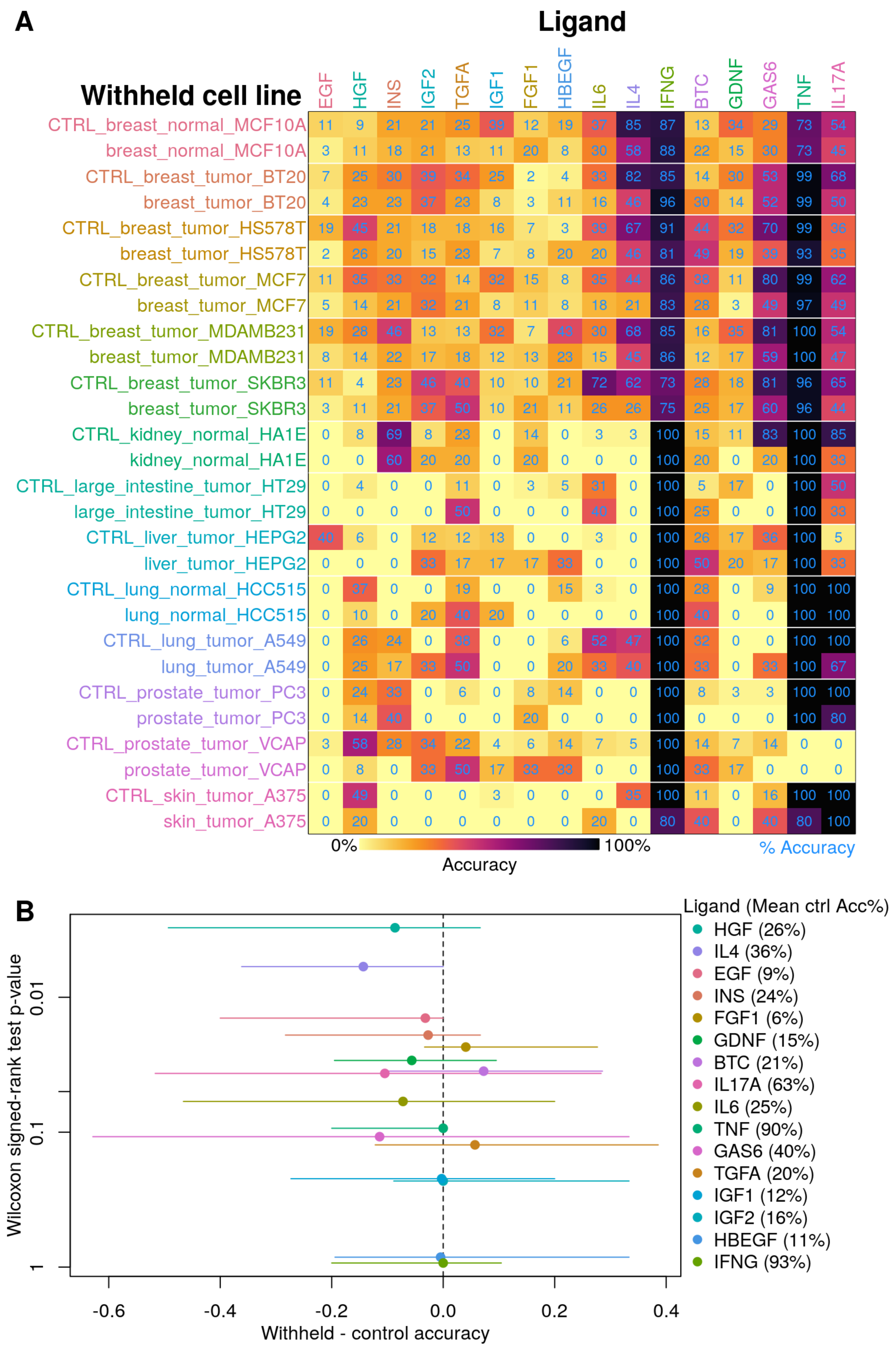

A
Ligand
Withheld cell line
EGF HGF INS IGF2 TGFA IGF1 FGF1 HBEGF IL6 IL4 IFNG BTC GDNF GAS6 TNF IL17A
CTRL_breast_normal_MCF10A 11 9 21 21 25 39 12 19 37 85 87 13 34 29 73 54
breast_normal_MCF10A 3 11 18 21 13 11 20 8 30 58 88 22 15 30 73 45
CTRL_breast_tumor_BT20 7 25 30 39 34 25 2 4 33 82 85 14 30 53 99 68
breast_tumor_BT20 4 23 23 37 23 8 3 11 16 46 96 30 14 52 99 50
CTRL_breast_tumor_HS578T 19 45 21 18 18 16 7 3 39 67 91 44 32 70 99 36
breast_tumor_HS578T 2 26 20 15 23 7 8 20 20 46 81 49 19 39 93 35
CTRL_breast_tumor_MCF7 11 35 33 32 14 32 15 8 35 44 86 38 11 80 99 62
breast_tumor_MCF7 5 14 21 32 21 8 11 8 18 21 83 28 3 49 97 49
CTRL_breast_tumor_MDAMB231 19 28 46 13 13 32 7 43 30 68 85 16 35 81 100 54
breast_tumor_MDAMB231 8 14 22 17 18 12 13 23 15 45 86 12 17 59 100 47
CTRL_breast_tumor_SKBR3 11 4 23 46 40 10 10 21 72 62 73 28 18 81 96 65
breast_tumor_SKBR3 3 11 21 37 50 10 21 11 26 26 75 25 17 60 96 44
CTRL_kidney_normal_HA1E 0 8 69 8 23 0 14 0 3 3 100 15 11 83 100 85
kidney_normal_HA1E 0 0 60 20 20 0 20 0 0 0 100 20 0 20 100 33
CTRL_large_intestine_tumor_HT29 0 4 0 0 11 0 3 5 31 0 100 5 17 0 100 50
large_intestine_tumor_HT29 0 0 0 0 50 0 0 0 40 0 100 25 0 0 100 33
CTRL_liver_tumor_HEPG2 40 6 0 12 12 13 0 0 3 0 100 26 17 36 100 5
liver_tumor_HEPG2 0 0 0 33 17 17 17 33 0 0 100 50 20 17 100 33
CTRL_lung_normal_HCC515 0 37 0 0 19 0 0 15 3 0 100 28 0 9 100 100
lung_normal_HCC515 0 10 0 20 40 20 0 0 0 0 100 40 0 0 100 100
CTRL_lung_tumor_A549 0 26 24 0 38 0 0 6 52 47 100 32 0 0 100 100
lung_tumor_A549 0 25 17 33 50 0 0 20 33 40 100 33 0 33 100 67
CTRL_prostate_tumor_PC3 0 24 33 0 6 0 8 14 0 0 100 8 3 3 100 100
prostate_tumor_PC3 0 14 40 0 0 0 20 0 0 0 100 0 0 0 100 80
CTRL_prostate_tumor_VCAP 3 58 28 34 22 4 6 14 7 5 100 14 7 14 0 0
prostate_tumor_VCAP 0 8 0 33 50 17 33 33 0 0 100 33 17 0 0 0
CTRL_skin_tumor_A375 0 49 0 0 0 3 0 0 0 35 100 11 0 16 100 100
skin_tumor_A375 0 20 0 0 0 0 0 0 20 0 80 40 0 40 80 100
0%
100%
Accuracy
% Accuracy
B
Wilcoxon signed-rank test p-value
0.01
0.1
1
-0.6 -0.4 -0.2 0.0 0.2 0.4
Withheld - control accuracy
Ligand (Mean ctrl Acc%)
HGF (26%)
IL4 (36%)
EGF (9%)
INS (24%)
FGF1 (6%)
GDNF (15%)
BTC (21%)
IL17A (63%)
IL6 (25%)
TNF (90%)
GAS6 (40%)
TGFA (20%)
IGF1 (12%)
IGF2 (16%)
HBEGF (11%)
IFNG (93%)

**Supplementary Figure 5. Accuracy in identifying which ligand treatment caused the changes in the transcriptome of a novel cell line.**

**a)** Random forest models were trained to identify the ligand treatment (columns) causing transcriptomic changes in a withheld cell line (rows), compared to performance when trained on all cell lines (leave-one-out testing, rows labeled ‘CTRL’). Darker colour represents higher accuracy, and number of samples per ligand and cell line is shown in blue.

**b)** Change in accuracy between predicting on a novel cell line versus the control model trained on all cell lines (x-axis - colours match ligand column names above). Lines represent range across cell lines with median indicated by a point on each line. Lines extending to the left of the dashed vertical line indicate that holding out a cell line reduces model performance, and vice versa for lines extending to the right of the dashed line. Wilcoxon signed-rank test was used to determine significance of pairwise change in accuracy (y-axis). Mean control accuracy is indicated in brackets in the legend to give context to the possible magnitude of AUPR change.

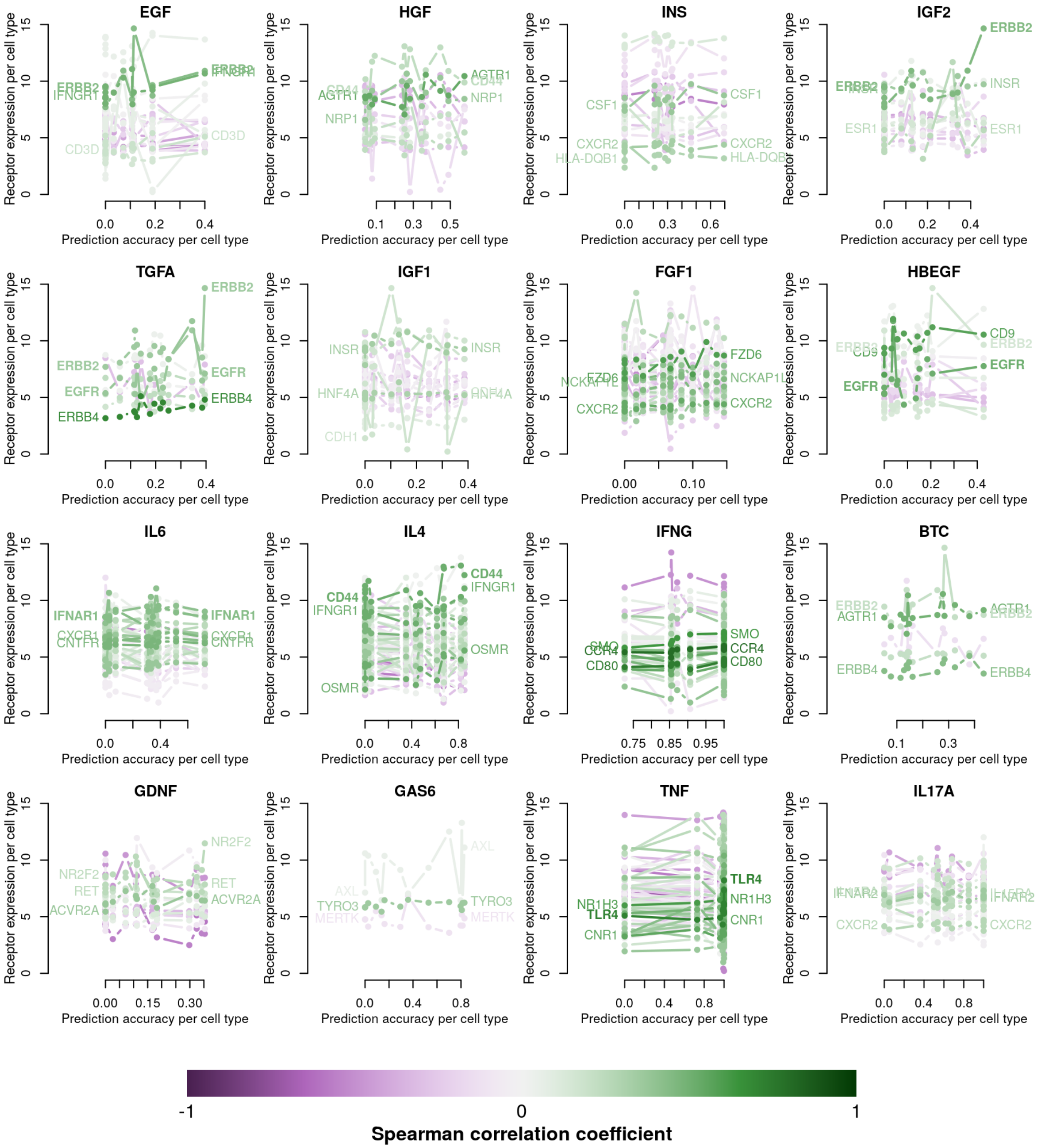


**Supplementary Figure 6. Cognate receptor gene expression does not sufficiently explain differences in transcriptional response to ligand perturbation.**

Cognate receptor expression for each ligand was compared to ligand classification accuracy per cell line. Lines for each receptor are coloured by the Spearman correlation coefficient between expression and accuracy per cell type (purple is negative, green is positive), and the three most correlated receptor genes are labeled.

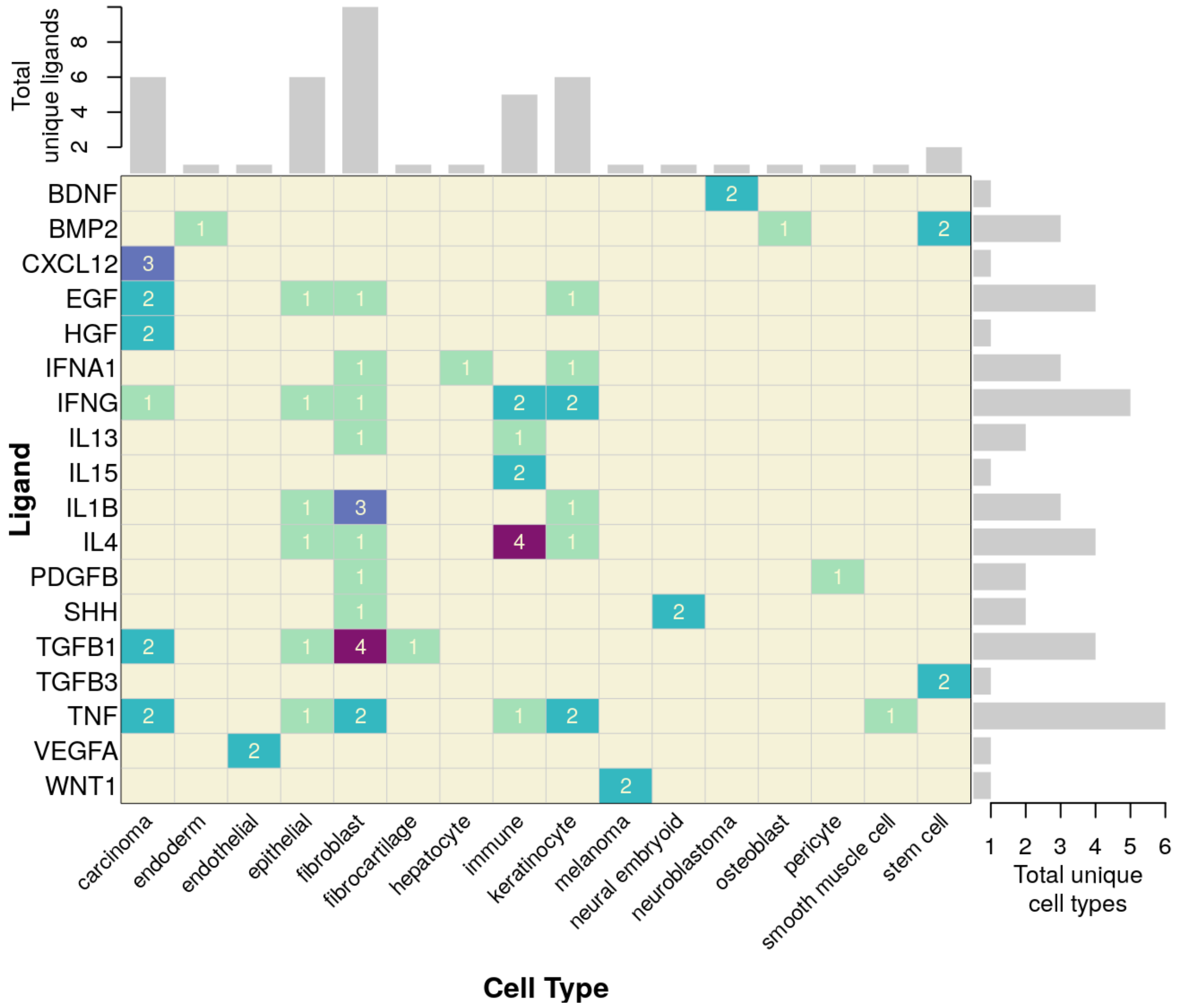


**Supplementary Figure 7. Distribution of experimental conditions from NicheNet-curated transcriptomics assays assessing change in gene expression caused by ligand treatment.**

The matrix indicates which ligands (rows) were assayed in which cell types (columns), with the number at each matrix element indicating how many transcriptional experiments from independent labs contained the same combination of ligand and cell type (darker colour reflecting higher count). The bar plots indicate row and column sums for the unique cell types in which a given ligand was assayed, and unique ligands assayed in a given cell type, respectively.

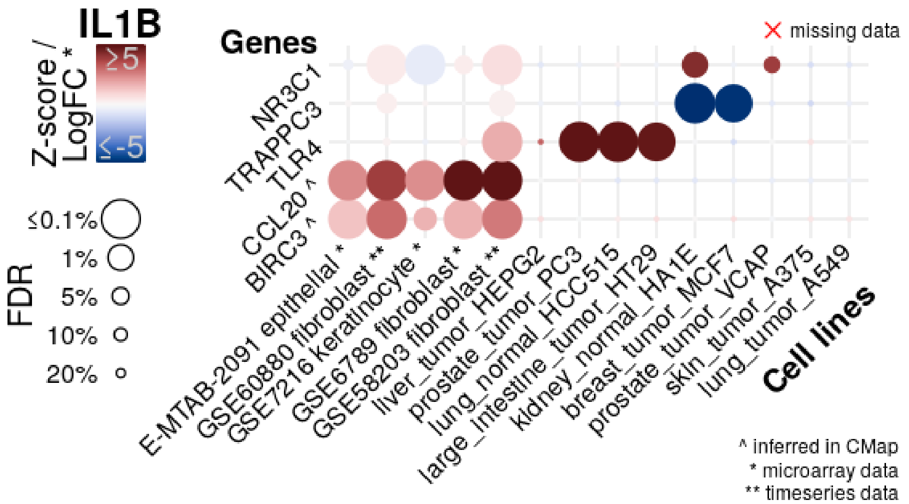
IL1B
Z-score / LogFC *
≥5
≤-5
FDR
≤0.1%
1%
5%
10%
20%
Genes
NR3C1
TRAPPC3
TLR4
CCL20 ^
BIRC3 ^
missing data
E-MTAB-2091 epithelial *
GSE60880 fibroblast **
GSE7216 keratinocyte *
GSE6789 fibroblast *
GSE58203 fibroblast **
liver_tumor_HEPG2
prostate_tumor_PC3
lung_normal_HCC515
large_intestine_tumor_HT29
kidney_normal_HA1E
breast_tumor_MCF7
prostate_tumor_VCAP
skin_tumor_A375
lung_tumor_A549
Cell lines
^ inferred in CMap
* microarray data
** timeseries data

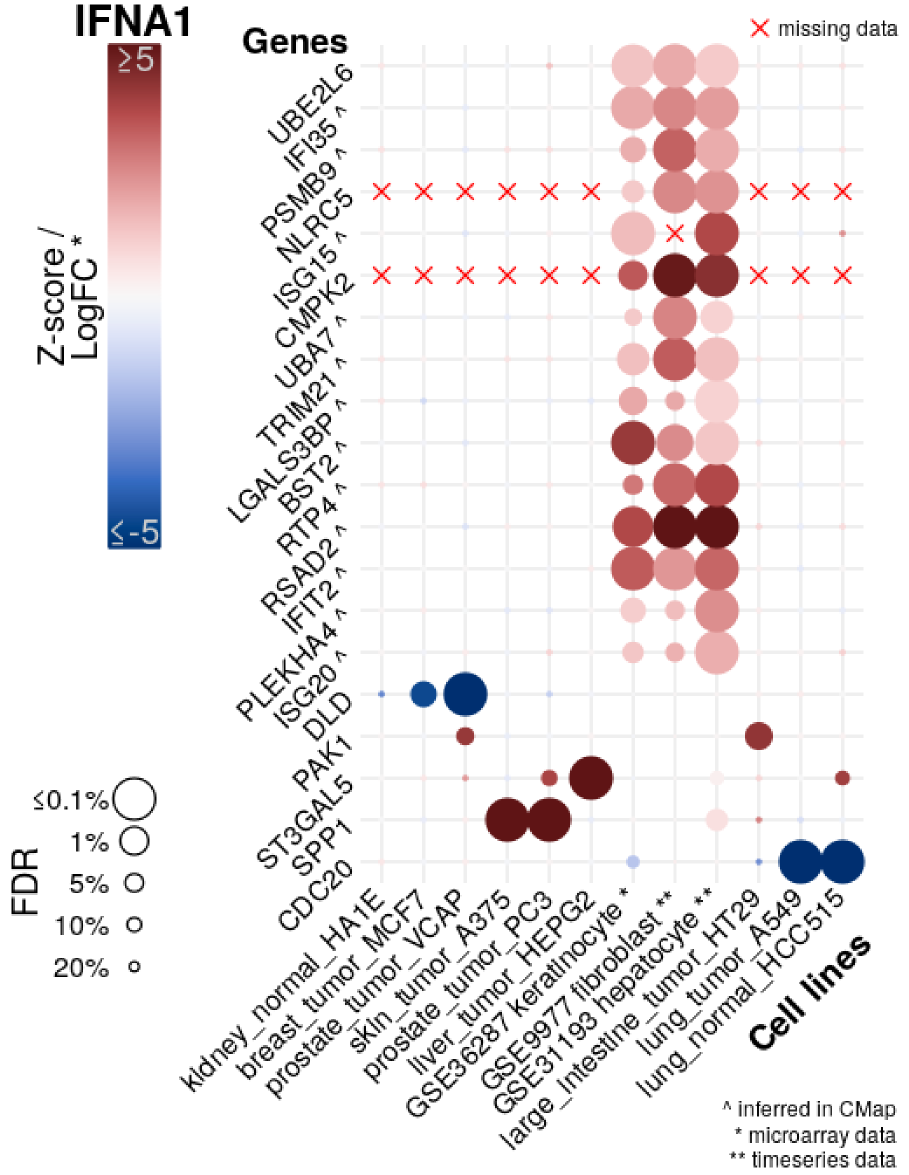
IFNA1
Z-score / LogFC *
≥5
≤-5
FDR
≤0.1%
1%
5%
10%
20%
Genes
UBE2L6
IFI35 ^
PSMB9 ^
NLRC5
ISG15 ^
CMPK2
UBA7 ^
TRIM21 ^
LGALS3BP ^
BST2 ^
RTP4 ^
RSAD2 ^
IFIT2 ^
PLEKHA4 ^
ISG20 ^
DLD
PAK1
ST3GAL5
SPP1
CDC20
missing data
kidney_normal_HA1E
breast_tumor_MCF7
prostate_tumor_VCAP
skin_tumor_A375
prostate_tumor_PC3
liver_tumor_HEPG2
GSE36287 keratinocyte *
GSE9977 fibroblast **
GSE31193 hepatocyte **
large_intestine_tumor_HT29
lung_tumor_A549
lung_normal_HCC515
Cell lines
^ inferred in CMap
* microarray data
** timeseries data

**Supplementary Figure 8. No general transcriptional response signature for IL1B or IFNA1 ligands.** Matrix plots showing mean change in gene expression per cell line upon IL1B or IFNA1 treatment for selected genes from both Connectivity Map and NicheNet-curated transcriptomics data. Cell lines are on each column, with NicheNet-curated transcriptomes indicated with *, and those from time series experiments where only magnitude of gene expression change was recorded with a **. Genes (in rows) were included if they were significantly differentially expressed (at 10% FDR) in at least two Connectivity Map cell lines or the maximum possible number of overlapping NicheNet-curated cell lines. Genes whose expression was inferred in Connectivity Map are indicated with ^. Genes whose expression was not reported in an experiment are indicated with a red x. Dot colour indicates difference in gene expression upon ligand treatment, measured as Connectivity Map Z-score and log2 fold-change from NicheNet-curated transcriptomes. Dot size indicates false discovery rate-corrected significance.

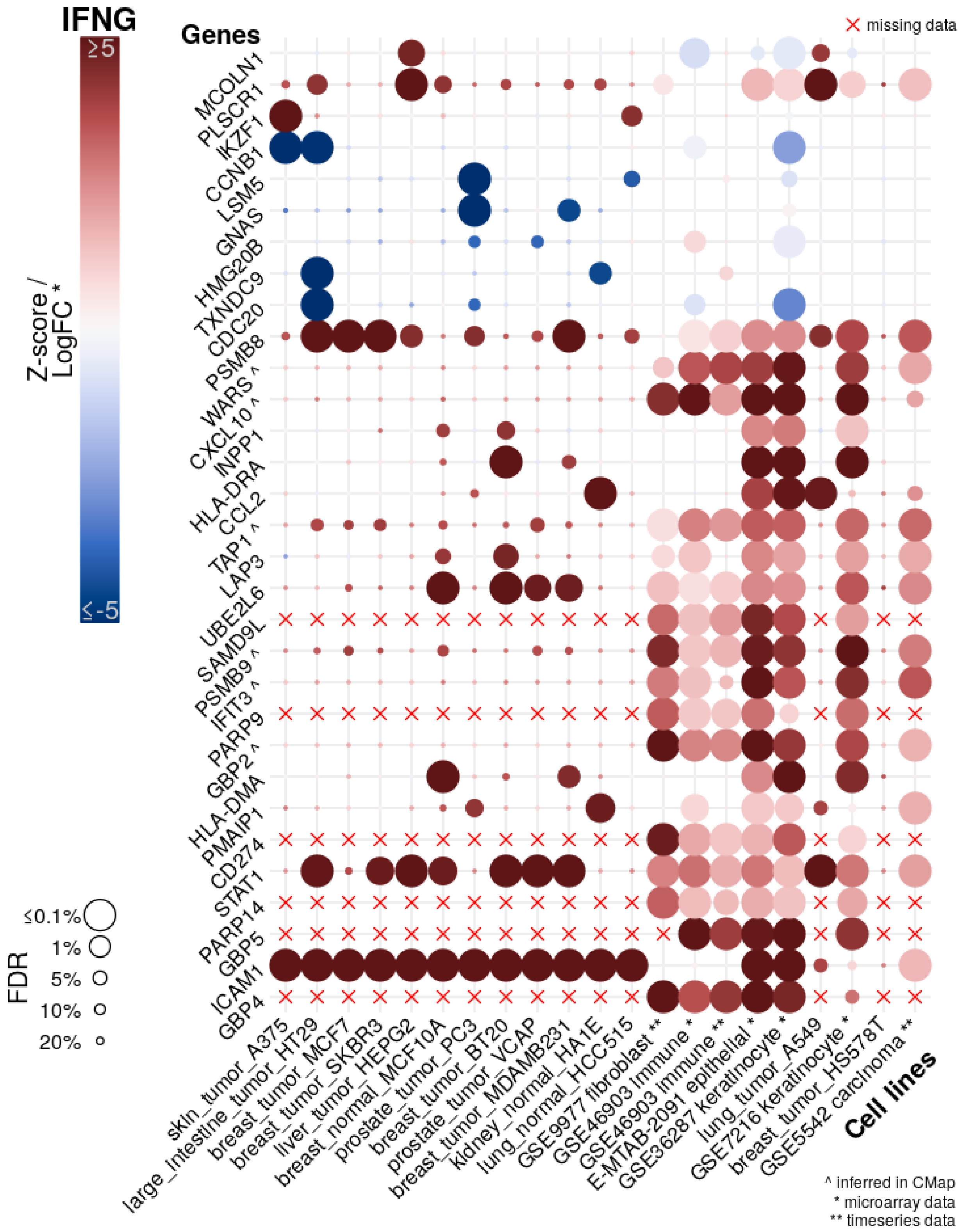


**Supplementary Figure 9. No general transcriptional response signature for IL1B or IFNG ligand.** A matrix plot showing mean change in gene expression per cell line upon IFNG treatment for selected genes from both Connectivity Map and

NicheNet-curated transcriptomics data. Cell lines are on each column, with NicheNet-curated transcriptomes indicated with *, and those from time series experiments where only magnitude of gene expression change was recorded with a **. Genes (in rows) were included if they were significantly differentially expressed (at 10% FDR) in at least two Connectivity Map cell lines or the maximum possible number of overlapping NicheNet-curated cell lines. Genes whose expression was inferred in Connectivity Map are indicated with ^. Genes whose expression was not reported in an experiment are indicated with a red x. Dot colour indicates difference in gene expression upon ligand treatment, measured as Connectivity Map Z-score and log2 fold-change from NicheNet-curated transcriptomes. Dot size indicates false discovery rate-corrected significance.

### Supplementary Tables

**Supplementary Table 1.** A comprehensive listing of novel methods and resources for cell-cell interaction prediction from transcriptomic data.

| Paper | Ligand-Receptor Source | Inference Method |
|---|---|---|
| Dynamic interaction networks in a hierarchically organized tissue (Kirouac *et al*, 2010) | Curated mostly from COPE (Ibelgaufts) | Ligand + receptor expression above threshold in microarray experiments of purified cell types. |
| Intercellular network structure and regulatory motifs in the human hematopoietic system (Qiao *et al*, 2014) | GO terms & manual curation for ligands, receptors. iRefWeb (Turner *et al*, 2010) for interactions. | Ligand + receptor expression above threshold in microarray experiments of purified cell types and deconvolved bulk RNAseq. |
| Transcriptome analysis of individual stromal cell populations identifies stroma-tumor crosstalk in mouse lung cancer model (Choi *et al*, 2015) | DLRP (Graeber & Eisenberg, 2001); HPRD (Keshava Prasad *et al*, 2009); KEGG (Kanehisa *et al*, 2016) | Ligand + receptor expression above threshold in bulk RNAseq experiments of purified cell types. |
| A draft network of ligand–receptor-mediated multicellular signalling in human (Ramilowski *et al*, 2015)<br>*Ligand-Receptor pairs from this paper are extensively referenced, referred to as the FANTOM5 database here.* | DLRP (Graeber & Eisenberg, 2001); IUPHAR (Harding *et al*, 2018); HPMR (Ben-Shlomo *et al*, 2003); curation for ligands, receptors.<br>HPRD (Keshava Prasad *et al*, 2009); STRING (Szklarczyk *et al*, 2015); curation for interactions. | Ligand + receptor expression above threshold in CAGE analysis of purified cell types (FANTOM5 dataset). |
| Proneurogenic ligands defined by modeling developing cortex growth factor communication networks (Yuzwa *et al*, 2016) | GO terms & manual curation for ligands, receptors. iRefWeb (Turner *et al*, 2010) for interactions. | Ligand + receptor expression above threshold in microarray experiments of purified cell types and deconvolved bulk. Cell surface proteomics for validation of receptor expression. |
| Social network architecture of human immune cells unveiled by quantitative proteomics (Rieckmann *et al*, 2017) | Secretome from experiments + STRING (Szklarczyk *et al*, 2015) | Proteomics from purified cell types. |

| | | |
|---|---|---|
| Single-cell transcriptomics of the human placenta: inferring the cell communication network of the maternal-fetal interface (Pavličev *et al*, 2017) | DLRP (Graeber & Eisenberg, 2001); IUPHAR (Harding et al. 2018) | Ligand + receptor expression above threshold in single-cell RNAseq experiments. |
| Extracting Intercellular Signaling Network of Cancer Tissues using Ligand-Receptor Expression Patterns from Whole-tumor and Single-cell Transcriptomes (Zhou *et al*, 2017) | FANTOM5 (Ramilowski *et al*, 2015) | Coexpression of ligand + receptor in TCGA samples. Ligand + receptor expression above threshold in single-cell RNAseq experiments. |
| Multilineage communication regulates human liver bud development from pluripotency (Camp *et al*, 2017) | FANTOM5 (Ramilowski *et al*, 2015) | Ligand + receptor expression above threshold in single-cell RNAseq experiments. |
| Single-Cell Transcriptional Profiling Reveals Cellular Diversity and Intercommunication in the Mouse Heart (Skelly *et al*, 2018) | FANTOM5 (Ramilowski *et al*, 2015) | Ligand + receptor expression above threshold in single-cell RNAseq experiments. |
| Lung Single-Cell Signaling Interaction Map Reveals Basophil Role in Macrophage Imprinting (Cohen *et al*, 2018) | FANTOM5 (Ramilowski *et al*, 2015) | Ligand + receptor expression above threshold in single-cell RNAseq experiments. |
| Analysis of Single-Cell RNA-Seq Identifies Cell-Cell Communication Associated with Tumor Characteristics (Kumar *et al*, 2018) | FANTOM5 (Ramilowski *et al*, 2015) | Ligand + receptor expression above threshold in single-cell RNAseq experiments. Scored by expression. |
| Single cell molecular alterations reveal target cells and pathways of concussive brain injury (Arneson *et al*, 2018) | UniProt “secreted” peptides as ligands. | Interaction score calculated by summing correlations between ligand and all genes (across cell types). Significance determined by permutation testing. |
| Single-cell transcriptomic profiling of the aging mouse brain (Ximerakis *et al*, 2019) | GO terms for ligands, receptors. Protein-protein interaction databases for links. Automated. | Ligand + receptor expression above threshold in single-cell RNAseq experiments. Ranked by differential expression. |
| Optimal-Transport Analysis of Single-Cell Gene Expression Identifies Developmental Trajectories in Reprogramming (Schiebinger *et al*, 2019) | GO terms for ligands, receptors. Protein-protein interaction databases for links. Curated. | Ligand + receptor interaction scored by positive coexpression in single-cell RNAseq experiments. |

| | | |
|---|---|---|
| Landscape of Intercellular Crosstalk in Healthy and NASH Liver Revealed by Single-Cell Secretome Gene Analysis (Xiong *et al*, 2019) | SPD (Chen *et al*, 2005) | Ligand + receptor expression above threshold in single-cell RNAseq experiments. |
| iTALK: an R Package to Characterize and Illustrate Intercellular Communication (Wang *et al*, 2019b) | DLRP (Graeber & Eisenberg, 2001); IUPHAR (Harding *et al*, 2018); FANTOM5 (Ramilowski *et al*, 2015) | Ligand + receptor expression above threshold in single-cell RNAseq experiments. |
| Dissecting intratumoral myeloid cell plasticity by single cell RNA-seq (Song *et al*, 2019) | IUPHAR (Harding *et al*, 2018); FANTOM5 (Ramilowski *et al*, 2015) | Ligand + receptor expression above threshold in single-cell RNAseq experiments. |
| Single-Cell Transcriptomics Analyses of Neural Stem Cell Heterogeneity and Contextual Plasticity in a Zebrafish Brain Model of Amyloid Toxicity (Cosacak *et al*, 2019) | FANTOM5 (Ramilowski *et al*, 2015) | Ligand + receptor expression above threshold in single-cell RNAseq experiments. |
| Adventitial Cell Atlas of wt (Wild Type) and ApoE (Apolipoprotein E)-Deficient Mice Defined by Single-Cell RNA Sequencing (Gu *et al*, 2019) | FANTOM5 (Ramilowski *et al*, 2015) | Ligand + receptor expression above threshold in single-cell RNAseq experiments. |
| Systematic expression analysis of ligand-receptor pairs reveals important cell-to-cell interactions inside glioma (Yuan *et al*, 2019) | FANTOM5 (Ramilowski *et al*, 2015) | Ligand + receptor expression above threshold in single-cell RNAseq experiments. Filtered for ligand-receptor pairs coexpressed in corresponding TCGA samples. |
| Single cell analysis of the developing mouse kidney provides deeper insight into marker gene expression and ligand-receptor crosstalk (Combes *et al*, 2019) | FANTOM5 (Ramilowski *et al*, 2015) | Ligand + receptor expression above threshold in single-cell RNAseq experiments. Networked weighted by gene expression, statistical model to find highly expressed interactions. |
| Single-cell expression profiling reveals dynamic flux of cardiac stromal, vascular and immune cells in health and injury (Farbehi *et al*, 2019) | FANTOM5 (Ramilowski *et al*, 2015) | Ligand + receptor expression above threshold in single-cell RNAseq experiments. Networked weighted by gene expression, statistical model to find highly expressed interactions. |
| Intra- and Inter-cellular Rewiring of the Human Colon during Ulcerative Colitis (Smillie *et al*, 2019) | FANTOM5 (Ramilowski *et al*, 2015) | Ligand + receptor expression above threshold in single-cell RNAseq experiments. Statistical model of cell-cell communication from number of ligand-receptor interactions per cell type pair. |

| | | |
|---|---|---|
| Single-cell reconstruction of the early maternal-fetal interface in humans (Vento-Tormo *et al*, 2018); CellPhoneDB: inferring cell-cell communication from combined expression of multi-subunit ligand-receptor complexes (Efremova *et al*, 2020) | IUPHAR (Harding *et al*, 2018); IMEx (Orchard *et al*, 2012); InnateDB (Breuer *et al*, 2013). | Statistical model for ligand-receptor enrichment from cell type pairs. Models complexes explicitly. |
| Cell lineage and communication network inference via optimization for single-cell transcriptomics (Wang *et al*, 2019a) | Curated selected pathways | Multilayer model of ligand-receptor, receptor-pathway, and gene regulatory networks. Probability of interaction is a function of ligand, receptor, and pathway gene expression. |
| Single-cell immune landscape of human atherosclerotic plaques (Fernandez *et al*, 2019) | FANTOM5 (Ramilowski *et al*, 2015) | Ligand + receptor expression above threshold in single-cell RNAseq experiments. Scored by expression. |
| Uncovering hypergraphs of cell-cell interaction from single cell RNA-sequencing data (Tsuyuzaki *et al*, 2019) | Organism-specific DBs using UniProt, HPRD (Keshava Prasad *et al*, 2009); STRING (Szklarczyk *et al*, 2015) | Ligand + receptor expression above threshold in single-cell RNAseq experiments. Modeled as a hypergraph, with cell types as nodes and L-R interactions are hyperedges, which allows for one/many-to-many interactions. |
| PyMINEr finds gene and autocrine-paracrine networks from human islet scRNA-seq (Tyler *et al*, 2019) | Extracellular domain or secreted GO terms; STRING (Szklarczyk *et al*, 2015) | Ligand + receptor expression above threshold in single-cell RNAseq experiments. |
| SingleCellSignalR: inference of intercellular networks from single-cell transcriptomics (Cabello-Aguilar *et al*, 2020) | FANTOM5 (Ramilowski *et al*, 2015); Reactome (Jassal *et al*, 2020); manual curation | Ligand + receptor expression above threshold in single-cell RNAseq experiments. Scored by normalized expression. |
| Single-cell transcriptome-based multilayer network biomarker for predicting prognosis and therapeutic response of gliomas (Zhang *et al*, 2020) | DLRP (Graeber & Eisenberg, 2001); HPRD (Keshava Prasad *et al*, 2009); KEGG (Kanehisa *et al*, 2016) | Multilayer model of ligand-receptor, receptor-pathway, and gene regulatory networks. Supported by prognostic power of EGFR signal in glioma. |
| NicheNet: modeling intercellular communication by linking ligands to target genes (Browaeys *et al*, 2020) | KEGG (Kanehisa *et al*, 2016); FANTOM5 (Ramilowski *et al*, 2015); IUPHAR (Harding *et al*, 2018); | Multilayer model of ligand-receptor, receptor-pathway, and gene regulatory networks. Weights trained on collection of ligand-perturbation gene expression data. |
| Predicting cell-to-cell communication networks using NATMI (Hou *et al*, 2020) | connectomeDB2020 (updated FANTOM5) (Hou *et al*, 2020) | Ligand + receptor expression above threshold in single-cell RNAseq experiments. Weighted by expression or specificity. |

| | | |
|---|---|---|
| FunRes: resolving tissue-specific functional cell states based on a cell–cell communication network model (Jung *et al*, 2020) | FANTOM5 (Ramilowski *et al*, 2015); OmniPath (Türei *et al*, 2016); Reactome (Jassal *et al*, 2020); MetaCore (Clarivate) | Multilayer model of ligand-receptor, receptor-pathway, and gene regulatory networks. Ligand-receptor interaction inferred by Markov chain from transcription factor expression. |
| Reconstruction of cell spatial organization from single-cell RNA sequencing data based on ligand-receptor mediated self-assembly (Ren *et al*, 2020) | FANTOM5 (Ramilowski *et al*, 2015) & curated list of immune cytokines / chemokines. | Ligand + receptor expression above threshold in single-cell RNAseq experiments. Scored by expression. |
| Pan-Cancer Analysis of Ligand–Receptor Cross-talk in the Tumor Microenvironment (Ghoshdastider *et al*, 2021) | FANTOM5 (Ramilowski *et al*, 2015) & curated list of immune checkpoint L-R pairs. | Ligand + receptor expression above threshold in single-cell RNAseq experiments. Direction of cell-cell signaling scored by relative expression of ligand/receptor between cell types. |
| CytoTalk: De novo construction of signal transduction networks using single-cell transcriptomic data (Hu *et al*, 2021) | FANTOM5 (Ramilowski *et al*, 2015) | Multilayer model of ligand-receptor and intracellular gene interaction networks. Prize-collecting Steiner forest algorithm identifies subnetwork with highest L-R correlation across cell types and intracellular cell type specificity. |
| Inference and analysis of cell-cell communication using CellChat (Jin *et al*, 2021) | KEGG (Kanehisa *et al*, 2016); STRING (Szklarczyk *et al*, 2015) & literature curation | Ligand + receptor significantly overexpressed in cell type from scRNAseq data. Statistical model includes agonist + antagonist cofactor expression when scoring interactions. |
| scConnect: a method for exploratory analysis of cell-cell communication based on single cell RNA sequencing data (Jakobsson *et al*, 2021) | IUPHAR (Harding *et al*, 2018) & manual association of molecular ligands with gene products required for synthesis | Ligand + receptor expression above threshold in single-cell RNAseq experiments. Scored by expression. For molecular ligands, score is the geometric mean of synthesis genes. |
| CrossTalkeR: Analysis and Visualisation of Ligand Receptor Networks (Nagai *et al*, 2021) | CellphoneDB (Efremova *et al*, 2020) | Summarizes CellPhoneDB output as a directed graph for the purpose of calculating network topology statistics. |
| CellTalkDB: a manually curated database of ligand–receptor interactions in humans and mice (Shao *et al*, 2021) | STRING (Szklarczyk *et al*, 2015); text mining of GO terms, NCBI gene annotation, PubMed abstracts | A ligand-receptor database meant to be used with existing cell-cell interaction algorithms. |
| Cellinker: a platform of ligand–receptor interactions for intercellular communication analysis (Zhang *et al*, 2021b) | Literature curation, & DLRP (Graeber & Eisenberg, 2001); HPRD (Keshava Prasad *et al*, 2009); IUPHAR (Harding *et al*, | Interaction database includes endogenous small molecules annotated as inorganic, metabolite, natural product, peptide and synthetic organic. Interaction database categorizes |

| | | |
|---|---|---|
| | 2018); CellphoneDB (Efremova *et al*, 2020) | cell-cell interaction types. Statistical model prioritizes L-R interactions specific to cell type pairs. |
| CellCall: integrating paired ligand–receptor and transcription factor activities for cell–cell communication (Zhang *et al*, 2021a) | connectomeDB2020 (Hou *et al*, 2020); CellTalkDB (Shao *et al*, 2021); Cellinker (Zhang *et al*, 2021b); CellChat (Jin *et al*, 2021) | Multilayer model of ligand-receptor, receptor-pathway, and gene regulatory networks. Scoring combines expression of L-R with expression of downstream regulon linked by KEGG (Kanehisa *et al*, 2016) and TF databases using enrichment statistics from GSEA (Subramanian *et al*, 2005). |
| Integrated intra- and intercellular signaling knowledge for multicellular omics analysis (Türei *et al*, 2021) | Curation from 26 external databases. | OmniPath now includes annotations for protein roles in intercellular signaling: surface/secreted ligand; receptor; ECM; adhesion; surface/secreted enzymes; transporter. |
| Deciphering cell–cell interactions and communication from gene expression (Armingol *et al*, 2021) | Collected from many published ligand-receptor lists | Indexed available ligand-receptor lists: https://github.com/LewisLabUCSD/Ligand-Receptor-Pairs |
